# Hepatic IRE1 Protects Against Septic Cardiac Failure

**DOI:** 10.1101/2023.03.14.532202

**Authors:** Mark Li, Roger R. Berton, Qingwen Qian, J. Alan Maschek, Biyi Chen, Elizabeth Barroso, Adam J. Rauckhorst, Thomas S. Griffith, Eric B. Taylor, James E. Cox, Vladimir P. Badovinac, Gökhan S. Hotamisligil, Long-Sheng Song, Ling Yang

## Abstract

Metabolic reprogramming in response to infection plays a critical role for septic survival. During a septic episode, the heart heavily relies on hepatic lipid particles to prevent heart damage and failure. Inositol-Requiring Enzyme 1 (IRE1) is the most conserved unfolded protein response (UPR) regulator that governs homeostasis of the endoplasmic reticulum (ER), the major site for lipid synthesis and processing. Here we show that hepatocyte IRE1 is indispensable for protecting against septic mortality in two different rodent models of experimental sepsis. The protective effect of hepatic IRE1 was not attributed to the inflammatory response since hepatic IRE1 deletion did not alter hepatic or systemic cytokine response. However, loss of IRE1 in the liver significantly augmented septic cardiac dysfunction in part due to a skewed immune-metabolic balance. Lipidomic and metabolomic analyses further revealed that loss of IRE1 in the liver compromised adaptive intrahepatic and circulating lipid reprogramming, including VLDL, in response to septic challenge. Furthermore, we identified that the protective effects against septic mortality are mediated by a non-canonical IRE1-dependent mechanism. Together, our study provides the first insight into how a disruption of hepatic ER-mediated lipid metabolic regulation promotes sepsis-associated cardiac immuno-metabolic imbalance.

## INTRODUCTION

Sepsis is a condition characterized by a life-threatening organ dysfunction caused by a dysregulated host response to infection (Singer et al., 2016), and it is recognized as the leading cause of death in the intensive care unit worldwide. Despite recent advances in the management of septic patients (Fleischmann et al., 2016; Motulsky, 1976), there are currently no effective FDA-approved agents to treat the patients.

Immunologic factors have been the primary target of translational and clinical investigation due to the presence of an overwhelming inflammatory cytokine response in septic patients and experimental animals (Chaudhry et al., 2013). However, numerous clinical trials that targeted various components of the innate immunity were not successful in improving septic outcomes. For example, clinical trials testing the effects of targeting IL-1(Opal et al., 1997) and TNF-a (Reinhart et al., 2001) did not yield benefits for sepsis-associated mortality. Additional trials that targeted toll-like receptor 4 (TLR-4) also did not provide any benefits for the patients (Opal et al., 2013). Therefore, our understanding of the pathophysiology of sepsis remains incomplete.

Aberrant adjustments of the systemic metabolic machinery can be detrimental for host survival. These include defective bioenergetics in different organs (Kwiatkowska, 1968) and abnormal levels of circulating metabolites (Langley et al., 2013). In addition to the aberrant immune response being a hallmark of sepsis, significant alterations in metabolism been observed in sepsis (Weis et al., 2017). Host’s response to sepsis requires extremely high amount of energy, which cannot be readily acquired by consuming food because septic individuals are unable or unwilling to eat due to anorexia (Murray and Murray, 1979; Wang et al., 2016; Weis et al., 2017). Therefore, sepsis creates a unique state of starvation response, which is intimately intertwined with concomitant systemic inflammation, and this septic immuno-metabolic interplay is not fully understood. Notably, recent studies suggest that certain plasma lipid species can be used as predictive markers of sepsis severity (Banoei et al., 2020; Ferrario et al., 2016; Khaliq et al., 2020; Lee et al., 2020). Indeed, supplementation of exogenous lipids, such as fish oil and ω-3 polyunsaturated fatty acids (PUFA), have beneficial effects on some clinical parameters in septic patients (Hall et al., 2015; Lu et al., 2017). However, it is unclear how to best modulate septic lipid reprograming in the clinic. Thereby, further research is needed to understand how lipids are dynamically regulated in the setting of sepsis.

The liver plays a central role in maintaining systemic homeostasis in sepsis (Strnad et al., 2017). In addition to controlling glucose and iron under septic conditions (Weis et al., 2017), the liver significantly contributes to lipid homeostasis (Alves-Bezerra and Cohen, 2017). For example, early clinical observations suggest the presence of a rapid influx of triglycerides primarily generated by the liver into circulation during a septic event (Harris et al., 2000). A recent study demonstrated that the liver serves as a critical source of lipids that are secreted in response to sepsis, which support cardiac function and host survival (Luan et al., 2019). However, it is unclear how this hepatic lipid response is mediated intracellularly on the molecular level. Hepatocytes, the major resident cell type in the liver, can sense an invading pathogen and secrete large quantities of lipid particles in the form of very-light-density-lipoproteins (VLDLs) as a response mechanism (Aspichueta et al., 2005). The primary site of lipid biosynthesis and modifications in the hepatocyte, including VLDL, is the endoplasmic reticulum (ER). Among many regulators of hepatic ER lipid homeostasis, Inositol-requiring enzyme 1 (IRE1) holds a central place (Fu et al., 2012; Hetz et al., 2020; Huang et al., 2019; Liu et al., 2019; Wang et al., 2018; Wang et al., 2012). However, the pathophysiological relevance of hepatic IRE1-dependent lipid regulation in sepsis is essentially unknown.

In this study, we investigated the role of hepatic ER-dependent lipid metabolism in sepsis. We show that hepatic IRE1 is necessary for protecting host against sepsis-associated mortality. We further demonstrate that hepatic non canonical IRE1-mediated systemic lipid metabolism is required for maintaining cardiac immuno-metabolic status in sepsis.

## RESULTS

### Hepatic IRE1 is necessary for survival in sepsis

To determine the effects of sepsis on the adaptive hepatic UPR, we measured the transcript and protein levels of IRE1 in the liver of wild-type (WT; C57BL6/J) male mice challenged with lipopolysaccharide (LPS) intraperitonially. As shown in Fig. 1A&B, LPS significantly reduced IRE1 both at the level of transcript and protein. Next, we sought to determine the necessity of hepatic IRE1 in the context of sepsis. To this end, we crossed mice with loxP sites flanking exons 20 and 21 of *Ern1* ( gene encodes IRE1; IRE1^FL/FL^) (Iwawaki et al., 2009) with transgenic mice harboring Cre recombinase under the control of albumin promoter (*Alb-Cre*) to achieve liver-specific IRE1 deletion (IRE1^ALB-CRE^). Next, we employed cecal ligation and puncture (CLP), a common surgical strategy to induce live poly-bacterial infection (Hubbard et al., 2005), in IRE1^FL/FL^ and IRE1^ALB-CRE^ mice. Deletion of IRE1 in the liver significantly attenuated the survival (∼25% survived compared to ∼60% in IRE1^FL/FL^ group) and worsened clinical score compared to IRE1^FL/FL^ mice after CLP (Fig.1C&D). To determine whether the increased mortality in IRE1^ALB-CRE^ mice was driven by defective resistance mechanisms (i.e., immune responses), we measured circulating cytokines as well as bacterial burden in different organs. However, we did not find significant differences in these parameters between IRE1^FL/FL^ and IRE1^ALB-CRE^ septic mice (Fig.1E&F). This finding suggests that hepatic IRE1 governs septic organ/tissue tolerance mechanisms (i.e., adaptation of organs in the presence of excessive inflammatory response).

**Figure 1.**
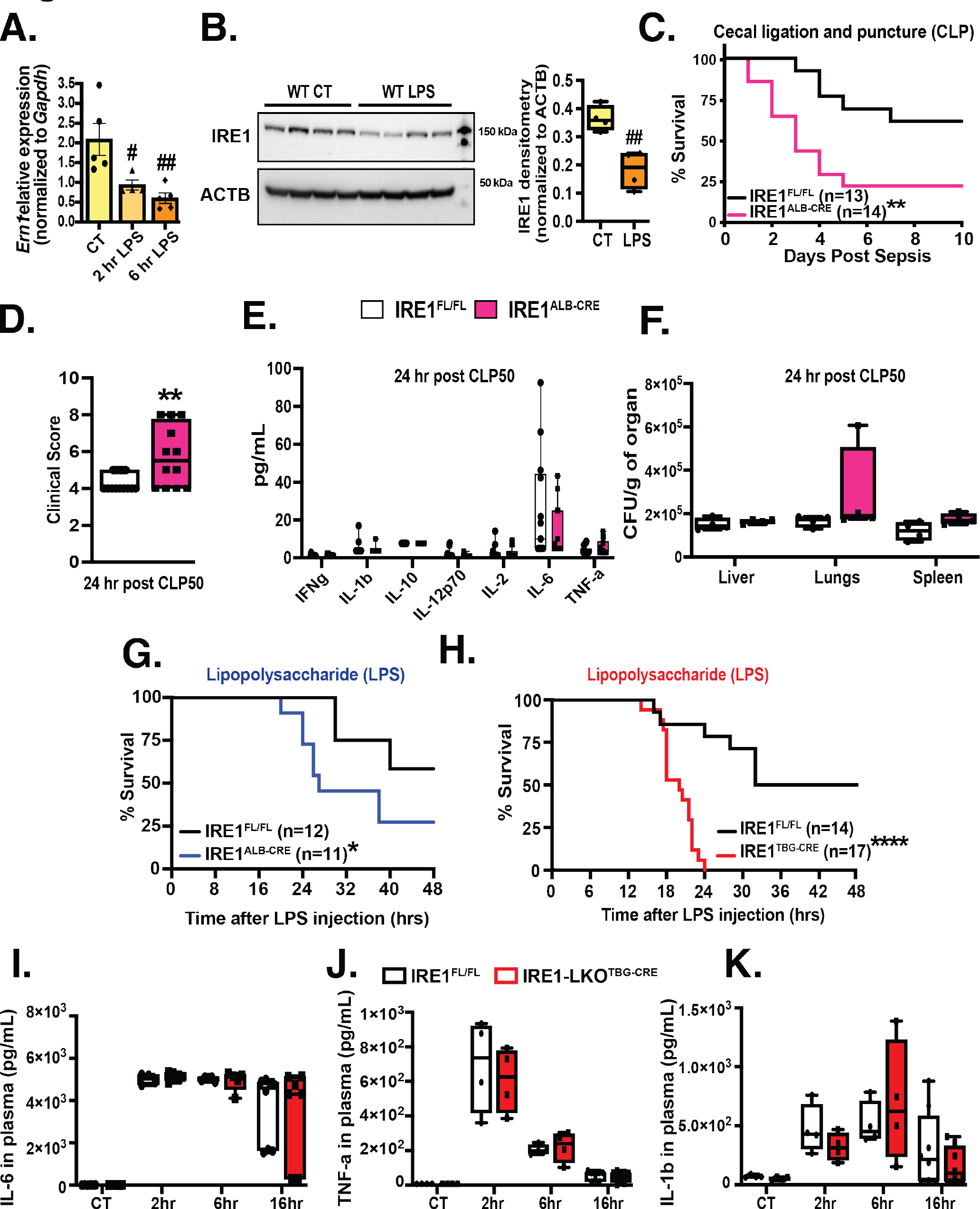
Hepatic IRE1 is necessary for survival in sepsis. **A**. *Ern1* transcript level measured by qRT-PCR in livers from WT animals challenged with LPS (12.5 mg/kg) for indicated time points and normalized to *Gapdh;* n = 5 age-matched male mice. **B**. Left: western blots of IRE1 and ACTB in livers from WT animals challenged with LPS (12.5 mg/kg) for 6 hours; Right: densitometry analysis of western blots; n = 4 age-matched male animals. **C**. Survival in IRE1^FL/FL^ and IRE1^ALB-CRE^ mice after cecal ligation and puncture (CLP); n = 13-14 age-matched male mice. **D**. Clinical score assessment in (**C**). **E.** Cytokines in plasma from IRE1^FL/FL^ and IRE1^ALB-CRE^ mice challenged with CLP for 24 hours and measured by Bio-Plex cytokine assay; n = 13-14 age-matched male mice. **F**. Bacterial load in indicated organs from IRE1^FL/FL^ and IRE1^ALB-CRE^ challenged with CLP as measured by qRT-PCR targeting the V4 region of the 16S rRNA gene and normalized to sample wet weight; n = 4 age-matched male mice. **G.** Survival in IRE1^FL/FL^ and IRE1^ALB-CRE^ mice challenged with LPS LD50; n = 11-12 age-matched male mice. **H.** Survival in IRE1^FL/FL^ and IRE1^TBG-CRE^ mice challenged with LPS LD50; n = 14-17 age-matched male mice. **I-K.** Plasma cytokines from IRE1^FL/FL^ and IRE1^TBG-CRE^ mice challenged with LPS LD50 as measured by ELISA kit; n = 8 age-matched male mice. Data are shown as means ± SEM. *Indicates genetic effects and # indicates CLP or LPS effects. Statistical significance was determined by Student’s t-test in (**A**), (**B**), & (**D**). Two-way ANOVA followed by Tukey’s multiple comparisons test was used in (**E**), (**F**) & (**I-K**). Log-rank (Mantel-Cox) test was used in (**C**), (**G**) & (**H**).

To determine how hepatic IRE1 orchestrates organ adaptation in response to inflammation, we next switched to intraperitoneal administration of LPS, which induces a septic event by mounting the host inflammatory response without introducing live bacteria. Consistent with the results from mice with CLP, mortality was increased in IRE1^ALB-CRE^ mice compared to that in IRE1^FL/FL^ mice in the setting of LPS-induced sepsis (Fig.1G). Because Alb-Cre causes a germline deletion and may potentially have developmental effects, we also determined whether deletion of IRE1 in the liver of adult animals recapitulates our previous findings. To this end, we transduced IRE1^FL/FL^ animals with GFP (IRE1^FL/FL^) or Cre under the expression adenovirus-associated virus serotype 8 with (AAV8) thyroid hormone-binding globulin (TBG) promoter (IRE1^TBG-CRE^) (SFig.1A). Two months after the viral transduction, the animals were subjected to LPS-induced sepsis. As shown in Fig. 1H, hepatic IRE1 deletion in adult mice augmented septic mortality, which is consistent with our observations in IRE1^ALB-CRE^ septic mice. Notably, we found comparable levels of circulatory cytokines between hepatic IRE1-deficient mice and their littermate controls (Fig.1I-K), suggesting that acute loss of IRE1 in adult animals does not affect resistance mechanisms. Together, these data demonstrate the necessary role of hepatic IRE1 for the host survival in sepsis in part through modulating tolerance mechanisms (i.e., adaptation of organs to excessive inflammatory stress).

### Hepatic IRE1 deficiency augments sepsis-associated cardiac outcomes by disrupting cardiac immuno-metabolic balance

Sepsis causes damage in multiple organs and tissues leading to septic mortality. Thus, we next sought to determine the cause of mortality of IRE1^TBG-CRE^ mice by profiling commonly assessed physiological parameters and organ damage markers in circulation. Hypothermia, hypoglycemia and hyperlactatemia are key features associated with sepsis severity (Van Wyngene et al., 2018). We found comparable circulating levels of lactate, glucose and body core temperature between LPS-treated IRE1^TBG-CRE^ and IRE1^FL/FL^ mice (Fig.2A-C). In addition, the levels of liver damage markers, namely alanine transaminase (ALT) and aspartate transaminase (AST), were comparable between IRE1^FL/FL^ and IRE1^TBG-CRE^ septic mice (Fig.2D&E). However, there was increased circulatory levels of hypoxanthine (Farthing et al., 2015) (Fig.2F) and cardiac troponin I (CTNI) (Fig.2G) in IRE1^TBG-CRE^ mice, suggesting that loss of IRE1 in the hepatocyte contributes to cardiac damage in sepsis. Thus, we further assessed cardiac function in these mice by performing trans-thoracic echocardiography. As shown in Fig.2H-J, loss of IRE1 in the liver significantly impaired left ventricular contractility evidenced by decreased wall motion, ejection fraction, stroke volume, and other parameters without affecting heart rate (SFig.1B&C). Together, these results indicate that hepatic IRE1 regulates cardiac function in response to a septic challenge.

**Figure 2.**
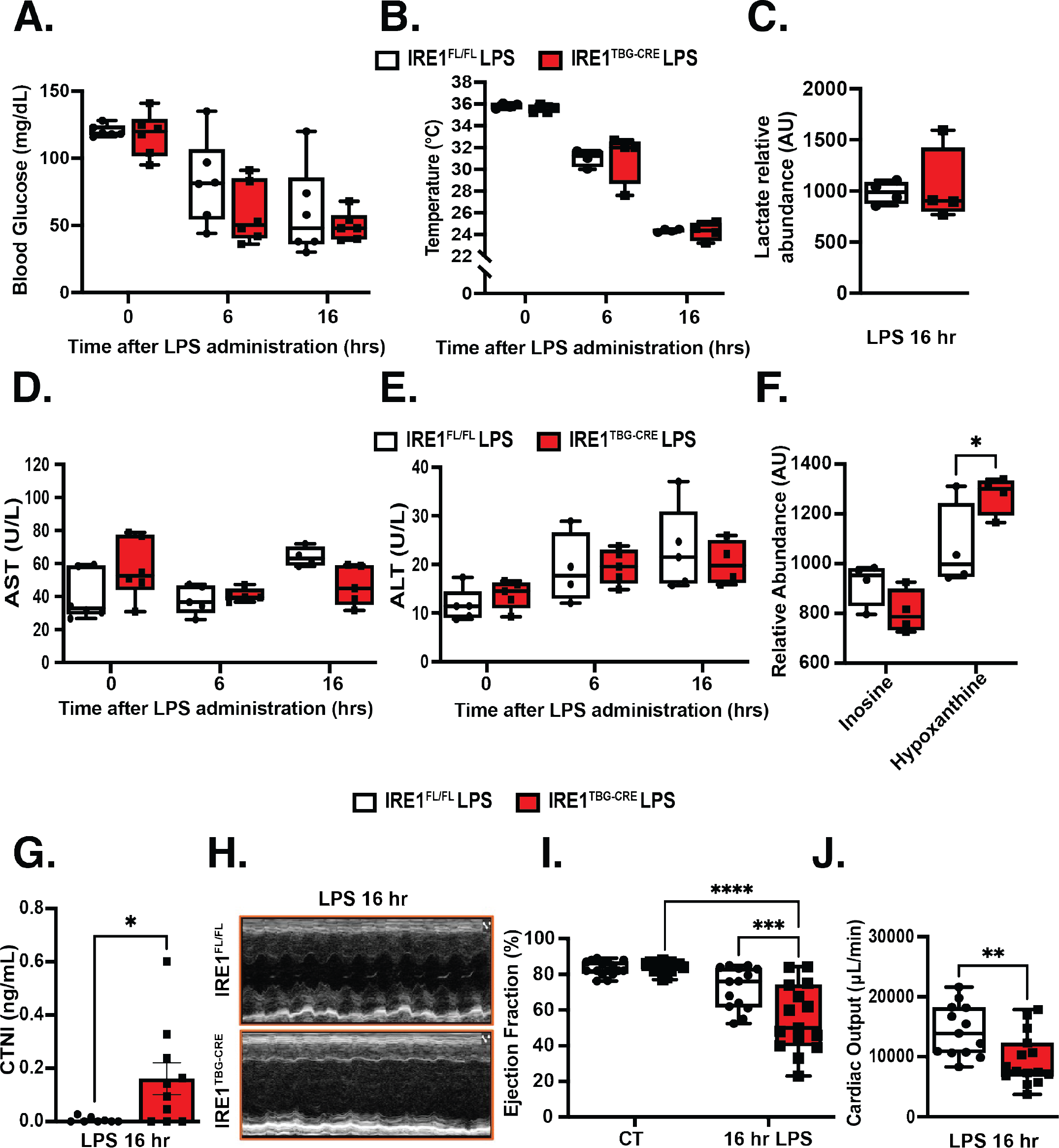
Hepatic IRE1 deficiency augments sepsis-associated cardiac outcomes. **A.** Blood glucose measured from tail vein in IRE1^FL/FL^ and IRE1^TBG-CRE^ mice challenged with LPS LD50 at indicated time points; n = 6 age-matched male mice. **B.** Body core temperature measured with a rectal probe in IRE1^FL/FL^ and IRE1^TBG-CRE^ mice challenged with LPS LD50 at indicated time points; n = 6 age-matched male mice. **C.** Lactate level in plasma from IRE1^FL/FL^ and IRE1^TBG-CRE^ mice challenged with LPS LD50 for 16 hours determined by targeted GC/MS; n = 4 age-matched male mice. **D&E.** ALT and AST levels in blood from IRE1^FL/FL^ and IRE1^TBG-CRE^ mice challenged with LPS LD50 at indicated time points; n = 6 age-matched male mice. **F.** Inosine and hypoxanthine levels in plasma from IRE1^FL/FL^ and IRE1^TBG-CRE^ mice challenged with LPS LD50 for 16 hours determined by targeted GC/MS; n = 4 age-matched male mice. **G.** CTNI in plasma from from IRE1^FL/FL^ and IRE1^TBG-CRE^ mice challenged with LPS LD50 for 16 hours measured by ELISA kit. **H.** Representative image of motion-mode (M-mode) in IRE1^FL/FL^ and IRE1^TBG-CRE^mice challenged with LPS LD50 for 16 hours measured with echocardiography. **I-J.** Ejection fraction and cardiac output in IRE1^FL/FL^ and IRE1^TBG-CRE^ mice challenged with LPS LD50 for 16 hours measured with echocardiography; n = 14-15 age-matched male mice. Data are shown as means ± SEM. *Indicates statistically significant genetic effects. Statistical significance was determined by Student’s t-test in (**C**), (**G**), & (**J**). Two-way ANOVA followed by Tukey’s multiple comparisons test was used in (**A-B**), (**D-F**) & (**I**).

To characterize the cellular and molecular impact of hepatic IRE1 on the septic heart, we carried out an RNA-Seq analysis in cardiac tissue isolated from IRE1^FL/FL^ and IRE1^TBG-CRE^ animals in the presence or absence of LPS septic challenge (SFig.2A-C). We focused on the 16-hour post-LPS time point, at which animals started to display mortality. As expected, cardiac inflammatory was significantly increased in mice treated with LPS (SFig.2D). Under basal conditions, deletion of hepatic IRE1 did not reprogram cardiac transcriptome (SFig.2C). In contrast, loss of hepatocyte IRE1 significantly altered the metabolic programming in the septic heart (Fig.3B), such as upregulating transcripts of amino acids across the membrane and downregulating transcripts of binding and uptake of ligands by scavenger receptors (Fig. 3A&B). Together, these findings indicate that loss of IRE1 in the liver disrupts septic-induced cardiac metabolic reprogramming.

**Figure 3.**
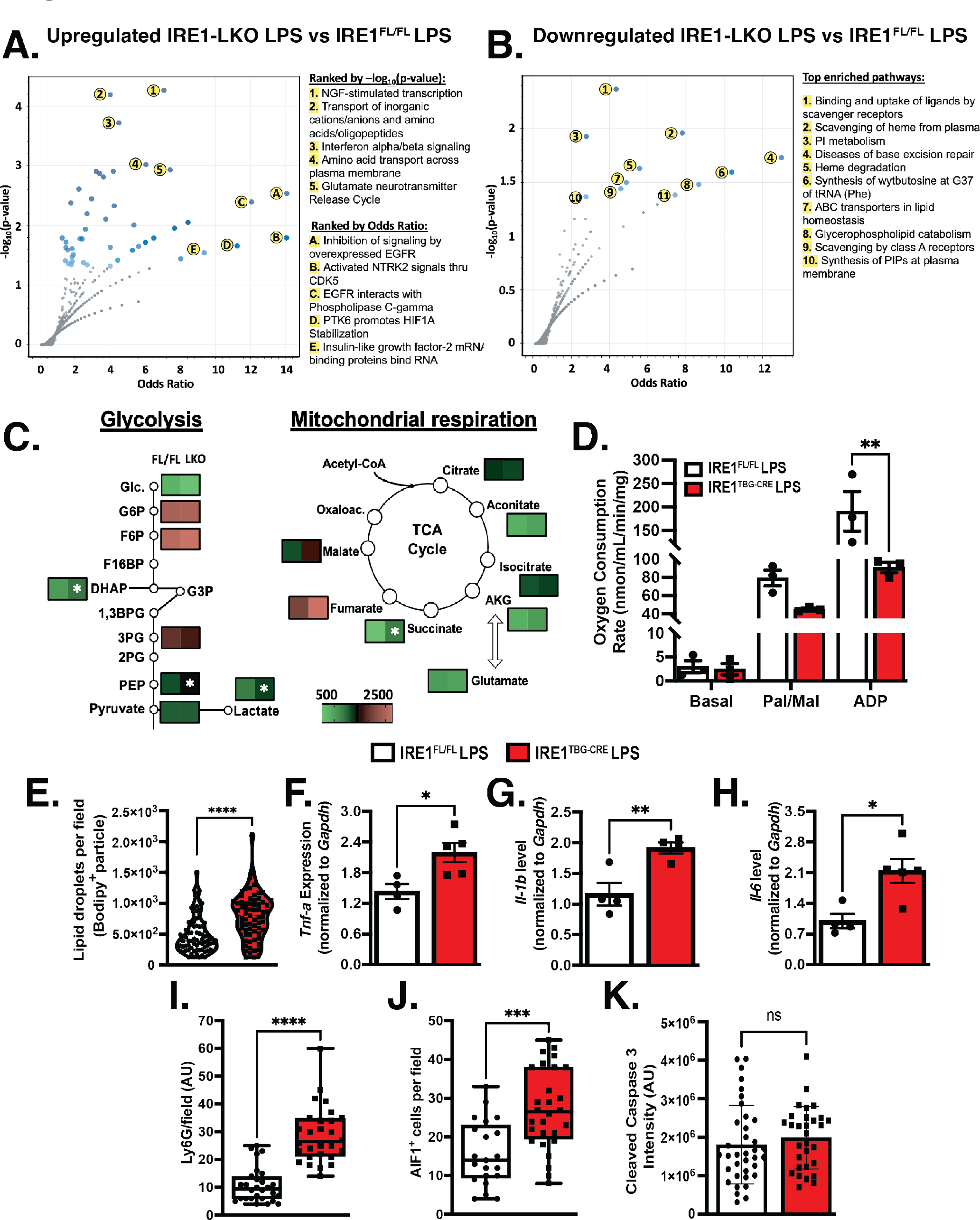
Hepatic IRE1 deficiency augments cardiac immuno-metabolic imbalance. **A-B.** RNA-Seq analysis performed in hearts from IRE1^FL/FL^ and IRE1^TBG-CRE^ mice challenged with LPS LD50 for 16 hours; REACTOME pathway analysis of upregulated (**A**) and downregulated (**B**) transcripts in IRE1^TBG-CRE^ versus IRE1^FL/FL^; n = 3 age-matched male mice. **C.** Steady-state GC/MS metabolomic analysis carried out in in hearts from IRE1^FL/FL^ and IRE1^TBG-CRE^ mice challenged with LPS LD50 for 16 hours; data are presented as fold change in IRE1^TBG-CRE^ versus IRE1^FL/FL^ hearts and Student’s t-test was performed for individual metabolites; n = 4 age-matched male mice. **D.** High-resolution respirometry (Oroboros instruments) in permeabilized cardiac fibers from IRE1^FL/FL^ and IRE1^TBG-CRE^ mice challenged with LPS LD50 for 16 hours; Pal/Mal – palmitoyl-carnitine/malate; ADP – adenosine diphosphate; n = 3 hearts from age- matched male mice. **E.** Quantification of lipid droplet intensity in the heart from IRE1^FL/FL^ and IRE1^TBG-CRE^ mice challenged with LPS LD50 for 16 hours as measured by Bodipy stain and acquired with confocal imaging; n = 5 age-matched male mice. **F-H.** Cytokine transcripts measured by qRT-PCR in the heart from IRE1^FL/FL^ and IRE1^TBG-CRE^ mice challenged with LPS LD50 for 16 hours and normalized to *Gapdh;* n =4-5 age-matched male mice. **I-K.** Quantification of immunofluorescence (**I&K**) or immunofluorescent particles (**J**) in hearts from IRE1^FL/FL^ and IRE1^TBG-CRE^ mice challenged with LPS LD50 for 16 hours acquired with confocal microscopy; n = 3-5 age-matched male mice. Data are shown as means ± SEM. *Indicates statistically significant genetic effects. Statistical significance was determined by Student’s t-test in (**E- K**). Two-way ANOVA followed by Tukey’s multiple comparisons test was used in (**D**).

To establish the metabolic impact of hepatic IRE1 deficiency on metabolic landscape in the heart in the context of sepsis, we next carried out a steady-state metabolomics analysis using hearts from IRE1^TBG-CRE^ and IRE1^FL/FL^ mice after 16 hours of LPS challenge (SFig2.E-G). Hepatic IRE1 deletion significantly reduced cardiac levels of DHAP, PEP, lactate, and succinate compared to IRE1^FL/FL^ mice (Fig. 3C). Of note, we did not observe significant differences in the levels of amino acids between the groups, except for elevated levels of alanine (SFig.2H). This finding may be explained by our RNA-Seq findings, whereby cardiac amino acid transport-associated transcripts were elevated in IRE1^TBG-CRE^ heart (Fig.3A). Additionally, we observed increased levels of intracardiac nucleotides in septic IRE1^TBG-CRE^ mice, potentially suggesting their active breakdown during septic event as a compensatory mechanism (SFig.2I).

At cellular level, previous studies indicate the presence of diminished mitochondrial respiration in septic heart, potentially exacerbating cardiomyopathy (Preau et al., 2021; Wasyluk et al., 2021). Therefore, to assess whether the disputed septic metabolic reprogramming in IRE1^TBG-CRE^ animals is associated with altered cardiac mitochondrial respiration, we performed high-resolution respirometry (Oroboros) analysis by using permeabilized cardiac fibers from mice subject to LPS stimulation (Mark Li, 2022; Schneider and Ayres, 2008). As shown in Fig.3D, the ADP-driven state 3 respiration was significantly reduced in cardiac fibers from mice with hepatic IRE1 deletion compared to control mice in the presence of palmitoyl-carnitine and malate. This finding was in line with the presence of altered levels of intermediate metabolites linked to the TCA cycle in the heart (Fig.3C) and increased abundance of lipid droplets in the ventricular wall of hearts from hepatic IRE1 deficient mice as measured by Bodipy staining (Fig.3E). Together, these results suggested that loss of IRE1 in the liver worsens cardiac mitochondrial activity in sepsis.

Another feature of septic heart is aberrant inflammatory response and increased immune cell infiltrates (Hollenberg and Singer, 2021; Silvestre-Roig et al., 2020; Zhang et al., 2023). We found that loss of hepatic IRE1 increased transcripts associated with interferon signaling in the heart compared to controls (Fig. 3A). We further assessed transcription levels of the cytokines associated with sepsis, namely *Il- 1b*, *Tnf-a*, and *Il-6*. As shown in Fig.3F-H, these proinflammatory transcripts were significantly elevated in the heart of IRE1^TBG-CRE^ but not in other organs (SFig.2J), suggesting a cardiac local inflammatory response. This finding was further supported by increased numbers of infiltrating macrophages and neutrophils in the ventricular wall of IRE1^TBG-CRE^ mice by immunohistochemistry (Fig.3I-J), but not apoptotic activation (Fig.3K). Together, these data indicate that deletion of hepatic IRE1 causes cardiac-specific dysfunction in part through regulating cardiac immuno-metabolic responses.

### Hepatic IRE1-mediated lipid reprogramming protects against septic cardiac damage

Previous studies implicated the role of IRE1 in regulating extrahepatic cholesterol and triglyceride (TG) levels after prolonged starvation and exposure to a high-fat diet (Wang et al., 2018; Wang et al., 2012; Yang et al., 2015). However, comprehensive and dynamic examination of hepatic IRE1 in regulating systemic lipidomic landscape in response to sepsis was not addressed in the past. To this end, we employed LC/MS steady-state lipidomic analysis to profile plasma lipids collected from IRE1^FL/FL^ and IRE1^TBG-CRE^ mice treated with vehicle or LPS at 2 and 6 hours of post injection (SFig.3). To enrich liver-derived factors, we specifically collected the blood from the suprahepatic inferior vena cava (SIVC) (Fig.4A). As expected, there was a dramatic increase in plasma TGs in IRE1^FL/FL^ mice in response to LPS, which was more apparent after 6 hours LPS administration (Fig.4C&D). Consistent with the previous studies (Claycomb et al., 1998; So et al., 2012; Wang et al., 2012), we observed a significant downregulation in major lipid groups in plasma from IRE1^TBG-CRE^ mice compared to IRE1^FL/FL^ throughout the tested conditions (Fig.4B-F & SFig.3E&G). In sepsis, adipose tissue undergoes uncontrolled lipolysis, thus liberating free fatty acids (FFAs) into circulation, which can enter peripheral organs including the liver (Azzu et al., 2020). However, our analysis did not detect any differences in circulating FFAs (Fig.4G), suggesting intact lipolysis from adipose tissue in mice with IRE1 deletion in the liver.

**Figure 4.**
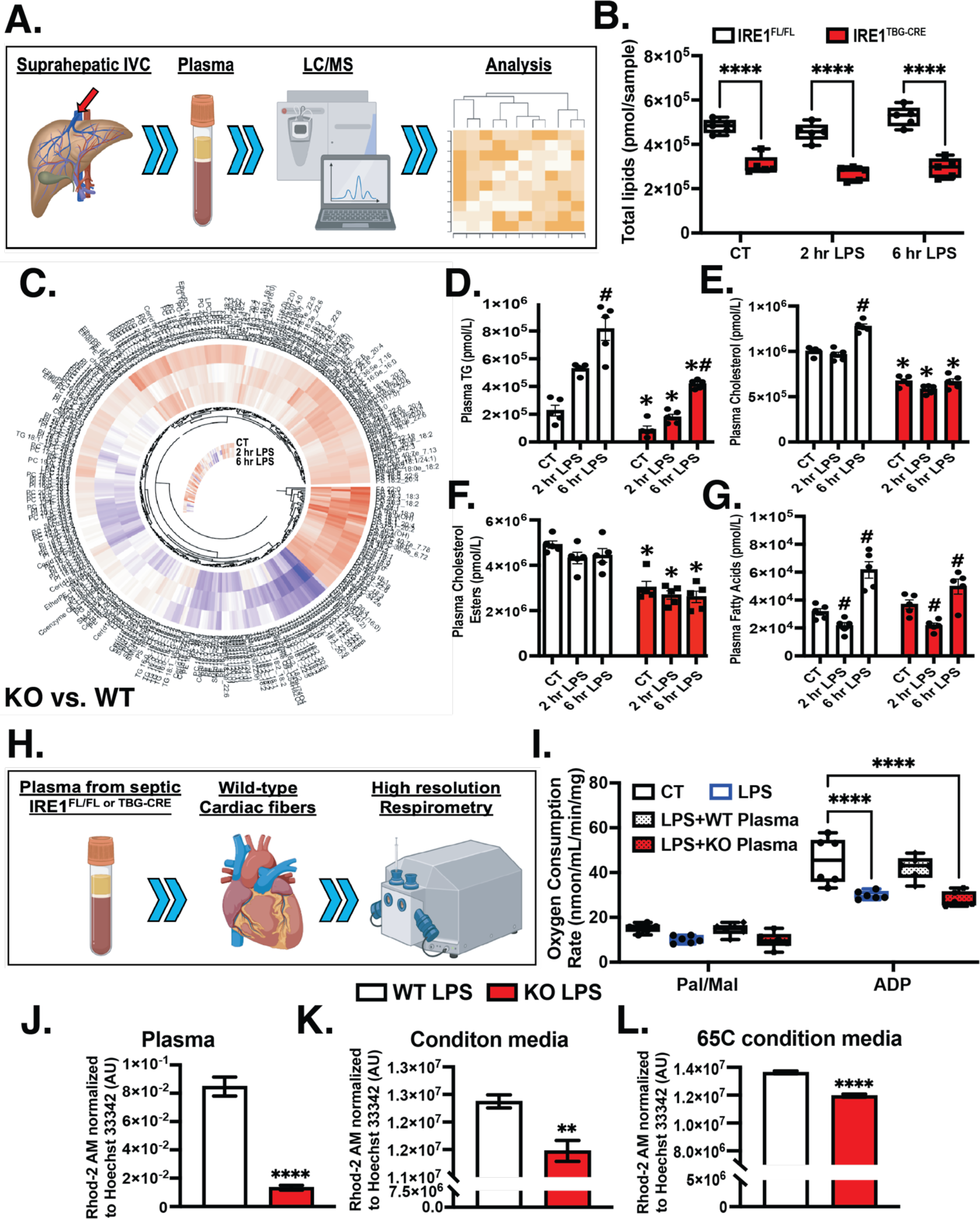
Hepatic IRE1-mediated lipid reprogramming protects against septic cardiac damage. **A.** Schematic representation of analyzing lipids in the plasma from IRE1^FL/FL^ and IRE1^TBG-CRE^ mice challenged with LPS at indicated time points. **B.** Total lipid abundance in the plasma from IRE1^FL/FL^ and IRE1^TBG-CRE^ mice at indicated time points; n = 5 age-matched male mice. **C.** Circular heatmap of detected lipid species presented as fold change IRE1^TBG-CRE^ vs IRE1^FL/FL^ for indicated conditions (outermost circle is CT, middle – 2 hr LPS, innermost – 6 hr LPS). **D-G**. Abundance of indicated lipid groups detected in the plasma from IRE1^FL/FL^ and IRE1^TBG-CRE^ mice at indicated time points; n = 5 age-matched male mice. **H.** Schematic representation of experimental setup for testing the effects of plasma on cardiac function. **I.** High- resolution respirometry (Oroboros instruments) in permeabilized cardiac fibers from WT mice challenged with LPS (2 µg/mL) for 3 hours in the presence of plasma from IRE1^FL/FL^ or IRE1^TBG-CRE^ septic mice; Pal/Mal – palmitoyl-carnitine/malate; ADP – adenosine diphosphate; n = 6 individual preparations from age-matched male mice. **J-L.** Intracellular calcium in HL-1 cells as measured by Rhod-2AM dye incubated with plasma from IRE1^FL/FL^ or IRE1^TBG-CRE^ mice (**J**), condition media from WT or IRE1 KO primary hepatocytes (**K**) or boiled condition media from WT or IRE1 KO primary hepatocytes (**L**); n = 3 individual experiments. Data are shown as means ± SEM. *Indicates genetic effects. Statistical significance was determined by Student’s t-test in (**J-L**). Two-way ANOVA followed by Tukey’s multiple comparisons test was used in (**B**), (**D-G**), & (**I**).

To determine whether hepatic IRE1-mediated protection for septic heart is attributed to plasma-associated molecules, we measured mitochondrial respiration in freshly permeabilized cardiac fibers from wildtype mice (WT, C57BL/6J) with or without a 3-hour treatment with LPS. In parallel, the LPS-treated fibers were also exposed to plasma collected from septic IRE1^FL/FL^ or IRE1^TBG-CRE^ animals (Fig.4H&I). LPS treatment significantly reduced WT cardiac fibers mitochondrial respiration, which was corrected by introducing plasma from IRE1^FL/FL^, but not plasma from IRE1^TBG-CRE^ mice (Fig. 4I).

We found that hepatic IRE1 deficiency augments sepsis-associated cardiac outcomes (Fig.2), but it was not clear whether plasma has direct effects on cardiomyocytes. Therefore, we next examined potential effects of plasma on cardiomyocyte calcium handling in a cardiomyocyte cell-line, HL-1 (Claycomb et al., 1998). As shown in Fig.4J, plasma from IRE1^TBG-CRE^ mice had suppressive effects on HL-1 calcium signaling compared to the IRE1^FL/FL^ group. Next, we asked whether these effects are mediated directly by hepatocyte-secreted factors. To this end, we measured calcium in HL-1 cells after an overnight incubation with condition media from WT or IRE1 KO primary hepatocytes challenged with LPS for 2 hours (Fig.4K) Finally, to address whether these effects are mediated by metabolites and not active enzymes present in condition media, we boiled it at 65℃ for 30 minutes before introducing it to HL-1 cells for calcium imaging. As shown in Fig.4L, boiled condition media isolated from IRE1-deficent hepatocytes reduced HL-1 calcium dynamics, suggesting metabolite/lipid-driven effects. Together, these data indicate that hepatocyte IRE1-mediated molecules have direct effects on cardiomyocyte function.

### RNase independent hepatic IRE1-VLDL axis is required for septic survival

To determine whether the reduction in plasma lipids by hepatocyte IRE1 deletion is due to alteration of intrahepatic lipid landscape, we carried out a steady-state untargeted lipidomic analysis of the liver in vehicle- and LPS-treated IRE1^FL/FL^ and IRE1^TBG-CRE^ mice at 2 and 6 hours after LPS challenge (Fig.5A & SFig.5A&B). Lipidomic analysis revealed overall accumulation of lipids due to IRE1 deletion, with elevated levels of TGs, cholesterol, and oxidized TGs being the prominent features (Fig. 5A&B & SFig.5C). This enriched intrahepatic lipid profile was further supported by measurement of lipid droplets in livers from IRE1^TBG-CRE^ mice compared to IRE1^FL/FL^ mice (Fig.5D&E, and SFig.4D&E).

**Figure 5.**
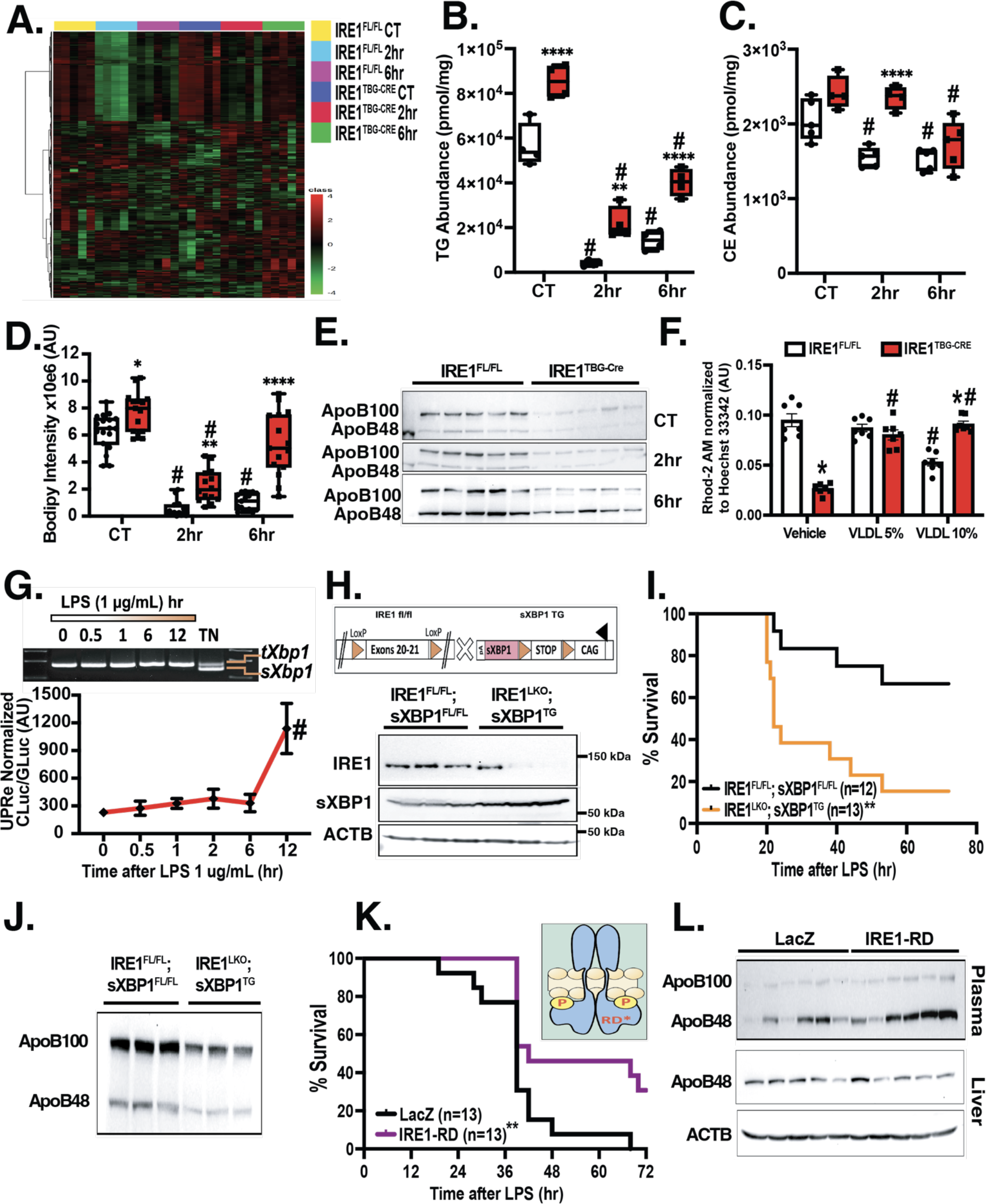
RNase independent role of hepatic IRE1 is required for septic survival. **A**. LC/MS lipidomic analysis carried out in livers from IRE1^FL/FL^ and IRE1^TBG-CRE^ mice challenged with LPS at indicated time points; n = 5 age-matched male mice. **B&C**. Intrahepatic levels of TG and cholesterol esters from IRE1^FL/FL^ and IRE1^TBG-CRE^ mice challenged with LPS at indicated time points measured with LC/MS lipidomics; n = 5 age-matched male mice. **D**. Lipid droplets as measured by Bodipy stain from IRE1^FL/FL^ and IRE1^TBG-CRE^ mice challenged with LPS at indicated time points; n = 5 age-matched male mice. **E.** Western blot analysis of Apob48/100 in plasma from IRE1^FL/FL^ and IRE1^TBG-CRE^ mice challenged with LPS at indicated time points; normalized to plasma volume; n = 5 age-matched male mice. **F.** Intracellular calcium signaling in HL-1 cells measured with Rhod-2AM in the presence of plasma from IRE1^FL/FL^ and IRE1^TBG-CRE^ mice and purified VLDL fraction from WT animals; n = 3 independent experiments. **G**. Top: spliced *Xbp1* measured by RT-PCR in primary hepatocytes challenged with LPS for indicated time points; Bottom: UPRe luciferase activity measured in primary hepatocytes challenged with LPS for indicated time points. **H**. Top: genetic strategy used to simultaneously delete and express IRE1 and sXBP1, respectively, in the liver; Bottom: western blot analysis of sXBP1, IRE1, and ACTB in the liver from the transgenic mice; n = 3 age-matched male mice. **I.** Survival of IRE1^FL/FL^; sXBP1^FL/FL^ and IRE1^LKO^; sXBP1^TG^ mice in response to LPS; n = 12-13 age-matched male mice. **J.** Western blot analysis of ApoB48/100 in plasma from IRE1^FL/FL^; sXBP1^FL/FL^ and IRE1^LKO^; sXBP1^TG^ mice challenged with LPS for 6 hours; n = 3 age-matched male mice. **K.** Survival of LacZ and IRE1^RD^ mice in response to LPS; n = 13 age-matched WT male mice. Data are shown as means ± SEM. *Indicates statistically significant genetic effects and # indicates effects of treatments. **L.** Western blot analysis of intrahepatic levels of ApoB and ACTB in WT mice transduced with adenovirus- LacZ or adenovirus-IRE1-RD for 2 weeks; n = 6 age-matched male mice. Data are shown as means ± SEM. One-way ANOVA with multiple test correction was used in (**G**); Two-way ANOVA followed by Tukey’s multiple comparisons test was used in (**B-D&F**); Log-rank (Mantel-Cox) test was used in (**I**)&(**K**).

Accumulation of triglycerides in the liver is attributed to multiple factors: increased releasing of FAs from adipose tissue, increased *de novo* synthesis of lipids, as well as impaired mitochondrial respiration and/or defective export of lipid-rich particles in the form of VLDLs (Cohen et al., 2011). We did not observe increased levels of FAs in the blood from IRE1^TBG-CRE^ mice (Fig.4G), suggesting a comparable lipolysis from adipose tissue in these mice. It is also known that *de novo* synthesis of lipids is suppressed in sepsis (Khaliq et al., 2020; Preau et al., 2021; Schulze et al., 2016; Trinder et al., 2019). Indeed, our attempts to carry out tracing experiments with acetyl-CoA into longer acyl chains did not yield a detectable signal (data not shown). To determine the hepatic lipid efflux, we measured levels of apolipoprotein B (ApoB; both ApoB48 and ApoB100) in plasma from IRE1^FL/FL^ and IRE1^TBG-CRE^ mice by western blot analysis. As shown in Fig.5E, hepatic IRE1 deletion decreased both basal as well and LPS-induced plasma ApoB compared to its littermate controls. This finding was associated with elevated intrahepatic ApoB levels (SFig.4F). Intracellularly, ApoB100 can be regulated transcriptionally and post-translationally (Haas et al., 2013). We next measured ApoB100 transcripts in liver from IRE1^FL/FL^ and IRE1^TBG-CRE^ mice, which did not reveal significant differences between the genotypes (SFig.4G). Coupled with our lipidomics analyses in the liver and circulation, these data suggest that hepatic IRE1 positively regulates secretion of ApoB particles from hepatocytes.

The secretion of ApoB requires its processing at the ER via lipidation, which ultimately forms apolipoprotein particles that are exported as VLDLs. We next aimed to determine whether reconstitution of VLDLs to plasma isolated from IRE1^TBG-CRE^ mice could rescue the cardiomyocyte calcium defect. To this end, we isolated VLDLs from WT animals injected with LPS for 2 hours (Li et al., 2018). We then measured calcium dynamics in cardiomyocytes (HL-1) supplemented with plasma from IRE1^FL/FL^ or IRE1^TBG-CRE^ septic animals in the absence (vehicle) or presence of WT VLDLs at 5% and 10% by volume. As shown in Fig.5F, VLDL is sufficient to rescue the defective hepatic IRE1-mediated calcium dysregulation in cardiomyocytes. Notably, 10% of VLDL had negative effects on HL-1 calcium signaling in the presence of IRE1^FL/FL^ plasma but not IRE1^TBG-CRE^ plasma, suggesting that excessive amounts of VLDL may lead to deleterious effects (Fig. 5F).

IRE1 possesses both kinase and RNase domains, which process a 26 base-pair fragment from XBP1, producing a spliced product (sXBP1), which can be translated and act as a transcription factor (Huang et al., 2019). Previously, Wang et al. implicated the crucial role of the IRE1-sXBP1 pathway in maintaining intrahepatic lipid homeostasis in response to prolonged starvation (So et al., 2012; Wang et al., 2012). To determine whether sXBP1 is involved in regulating the hepatic response to sepsis, we first assessed translocation of sXBP1 into the nucleus in the whole liver and found elevated levels of sXBP1 in the nuclear fraction (SFig.4H&I). Given that sepsis alters hepatic immune cells and other cell types composition (Martinon et al., 2010; Strnad et al., 2017), we next determined the cell autonomous role of hepatocyte XBP1 in response to sepsis. To this end, we isolated primary hepatocytes from WT animals and assessed the splicing event of XBP1 under septic conditions. To our surprise, treating the cells with LPS was not sufficient to induce the splicing event of XBP1 (Fig.5G, top panel), which is consistent with unaltered sXBP1 activity in response to LPS as determined by the UPRe luciferase reporter assay (Fu et al., 2015) (Fig.5G; bottom panel). Furthermore, to test whether reconstitution of sXBP1 into the liver could rescue septic mortality of hepatic IRE1-deficient animals, we crossed IRE1^FL/FL^ mice with transgenic mice with tissue-specific overexpressing sXBP1 (sXBP1^FL-STOP-FL^) (Zhang et al., 2021). These animals were then transduced with AAV8-TBG-iCre to delete IRE1 and simultaneously reconstitute sXBP1 in the liver (IRE1^TBG-CRE^; sXBP1^LTG^) (Fig.5H). Restoration of hepatic sXBP1 neither improved septic survival (Fig.5I) nor increased plasma ApoB levels (Fig.5J) in mice with lethal dose 50 (LD50) of LPS. These results suggest that sXBP1-dependent signaling might be dispensable in the context of IRE1-mediated metabolic adaptation in sepsis.

Lastly, to define whether IRE1’s RNase domain provides protection against sepsis-associated mortality, we overexpressed either a mutant form of IRE1 with dead RNase (IRE1^RD^) or control construct in the liver of WT male mice by adenovirus-mediated gene delivery (SFig.4J). Two weeks after the virus injection, the mice were subjected to sepsis by LPS injection. As shown in Fig.5K, overexpression of IRE1^RD^ in the liver significantly improved septic survival compared to control group. Furthermore, at the cellular level, this was associated with an increase in plasma levels of ApoB and a slight reduction of intracellular ApoB (SFig.5L). Together, our data indicate the pathophysiological role of RNase domain independent hepatic IRE1 for protecting mice against septic mortality.

## DISCUSSION

Sepsis elicits major spatio-temporal changes throughout the body, and failure to properly adapt results in a highly complex and life-threatening condition (Evans, 2018). Therefore, identifying roles of different components of response mechanisms requires a multi-organ approach with consideration of temporal progression of sepsis. In this study, we investigated the regulatory role of liver-to-heart crosstalk during early stages of a septic event. We identified hepatic IRE1 as the major regulator of liver-mediated lipid response that has significant impact on heart function and host survival.

A main feature of sepsis during the early phase is effective mounting of the inflammatory response mediated by cells of the innate immunity (Schneider and Ayres, 2008). In addition, sepsis is accompanied by both behavioral and cellular metabolic adaptations that are characterized by anorexia and starvation-like rewiring in metabolic organs, respectively (Medzhitov et al., 2012; Soares et al., 2014). Together, these responses aim to eliminate an invading pathogen and adapt according to host’s needs in order to maintain immuno-metabolic homeostasis. The liver holds a central place in the context of sepsis, as it orchestrates both the immune (e.g., acute phase reaction proteins.) and metabolic outputs (e.g., lipids) (Strnad et al., 2017). Particularly, hepatic ER is the main site that regulates these events by synthesizing and modifying proteins and lipids (Fu et al., 2012). Upon various stressors, including the presence of a pathogen, the ER undergoes significant structural and functional changes that, if uncontrolled, can lead to deleterious effects (Hetz et al., 2020; Hotamisligil, 2010). Consequently, cells induce the UPR in order to regain homeostasis in the ER (Hetz et al., 2020; Hotamisligil, 2010). Mounting evidence demonstrates that both inflammatory and metabolic cues can activate the most conserved branch of the UPR, IRE1, in different cell types (Hetz et al., 2020; Hotamisligil, 2010). For example, it was shown that LPS-mediated TLR signaling activates IRE1 in macrophages and adipocytes leading to cytokine production and lipolysis, respectively (Foley et al., 2021; Martinon et al., 2010). In addition, metabolic signals such as prolonged starvation, activates IRE1-XBP1-mediated processing and secretion of lipid particles in hepatocytes (So et al., 2012; Wang et al., 2012). However, precise mechanisms underlying the role of IRE1 in integrating immuno-metabolic signaling in sepsis, remain incompletely understood.

Lower levels of plasma lipids are associated with worse clinical outcomes in patients with sepsis (Lee et al., 2015; Maile et al., 2020). It has been demonstrated that, during the early phase of sepsis, both humans and laboratory animals display elevated levels of circulating lipids mostly due to liver- dependent lipidation and secretion of VLDLs (Harris et al., 2000). Indeed, pre-incubation of VLDLs with LPS before inducing endotoxemia or administration of triglyceride-rich lipoproteins after endotoxemia, protects mice and rats (Harris et al., 1990; Read et al., 1995) respectively, against sepsis-associated mortality. Notably, the occurrence or worsening of liver function during sepsis puts patients at higher risk of mortality (Nesseler et al., 2016). This evidence suggests that normal liver function, partly due to its ability to secrete VLDLs, may be crucial for providing protection against sepsis. In our study, we found the hepatic IRE1-dependent lipid/VLDL reprograming has protective role in heart function and host survival. A recent work identified GDF15-mediated liver-heart interaction in the setting of sepsis (Luan et al., 2019). Of note, Zhang et al. showed that Gdf15 can be directly regulated by sXBP1 in the liver during fasting (Zhang et al., 2018). However, we did not observe significant changes in the transcript of *Gdf15* between IRE1^FL/FL^ and IRE1^TBG-CRE^ animals (SFig.4K). Furthermore, previous studies demonstrate that hepatic IRE1 deficiency promote hypolipidemia in the context of starvation (So et al., 2012; Wang et al., 2012). Although sepsis yields a starvation-like response, it should be noted that, together with concomitant systemic inflammation, it may transmit signals to hepatic ER in a unique fashion, potentially including inputs from both DAMPs and PAMPs and various metabolic messengers. Mechanistically, it is well recognized that IRE1-induced sXBP1 transcriptional programs regulate hepatic metabolic homeostasis (Huang et al., 2019). However, accumulating evidence reveals noncanonical metabolic roles of IRE1 that occur in part through interactions with other proteins (Hetz et al., 2020; Huang et al., 2019). Although prolonged starvation requires hepatic IRE1-XBP1 transcriptional program for VLDL processing (So et al., 2012; Wang et al., 2012), we found that mice with hepatic IRE1 regulates secretion of ApoB in the context of sepsis is XBP1- and IRE1 RNase- activity-independent. (Fig.5I&L). Therefore, further research is necessary to establish context-dependent spatial-temporal localization of IRE1 and IRE1 interactome in hepatocyte. Moreover, although we found that VLDL is sufficient to correct the calcium defect in septic cardiomyocyte (Fig.5F), it is unclear how the hepatic IRE1-controled VLDLs provide cardiac protection in sepsis. Due to uncontrolled lipolysis of adipose tissue that occurs in sepsis, it is plausible to speculate that VLDLs may provide signaling molecules rather than being utilized as energy substrates in the heart for survival in sepsis.

In summary, our study demonstrates the metabolic role of hepatic IRE1 in sepsis-related cardiac dysfunction and provides novel insights into lipids mediated the liver-to-heart communication in the context of infection-induced cardiac dysfunction.

## Supporting information

Manuscript

## ACKNOWLEDGMENTS

We are grateful to Dr. Tiangang Li (OU College of Medicine, Oklahoma) and Dr. Yanqiao Zhang (NEOMED, Ohio) for technical supports. We thank Kathy Zimmerman and Alissa Bosko at the University of Iowa Cardiology Mouse Phenotyping Core Laboratory for performing transthoracic echocardiography. We also thank Drs. Vitor Lira, Julien Sebag and Brandon Davis at the University of Iowa for intellectual supports. Metabolomics analysis was performed at the Metabolomics Core Facility at the University of Utah. Mass spectrometry equipment was obtained through NCRR Shared Instrumentation Grant 1S10OD016232-01, 1S10OD018210-01A1 and 1S10OD021505-01.

## AUTHOR CONTRIBUTIONS

L.Y. & M.L. designed the study and wrote the manuscript. ML., R.B., Q.Q., B.C., E.B., A.J.R., & J.A.M. performed the experiments. A.J.R., E.B.T., T.S.G., J.C., V.B., G.S.H.& L.S. provided critical reagents, shared their expertise, constantly provided intellectual input, and revised the manuscript. L.Y. conceived and supervised the study. M.L., R.B., V.B., LS.S., and L.Y. are supported by the Collaborative Sciences Award (959706) from the American Heart Association. V.P.B. is supported by R35GM134880 and a University of Iowa Distinguished Scholar. R.R.B. is supported by T32AI007485.

## DECLARATION OF INTEREST

No conflict of interest is reported.

## METHODS

### Mouse models and experimental sepsis models

Animal experiments were carried out in accordance with the Guide for the Care and Use of Laboratory Animals (National Institutes of Health publication 85–23, revised 1996) and were approved by the Institutional Animal Care and Use Committee at the University of Iowa. All mice were bred and maintained under specific pathogen free conditions, and temperature was maintained at 22C. Mice were kept on a 12-hr light-dark cycle and fed a regular diet (Teklad global diet, 7913). *Ire1^fl^*mice were from Dr. Takao Iwawaki at the Gunma University (Japan). C57BL/6J mice were purchased from Jackson laboratory. *sXbp1^fl^* mice (*Hprt^tm1(fl-STEP-fl-sXbp1)Hota^*) were generated by genOway (France) (Zhang et al., 2021).

### Cecal-ligation and puncture (clp)-induced sepsis

Polymicrobial sepsis was induced by cecal ligation and puncture (CLP) (Berton et al., 2022; Sjaastad et al., 2020). Briefly, mice were anesthetized with ketamine/xylazine, abdomen was disinfected with Betadine (Purdue Products), and a midline incision was made. Subsequently, the distal third of the cecum was ligated with Perma-Hand Silk (Ethicon), punctured once using a 25-gauge needle, and a small amount of fecal matter was extruded out of puncture. The cecum was then returned to the abdomen, the peritoneum was closed with 641G Perma-Hand Silk (Ethicon). Bupivacaine (Hospira) was then administered at incision site, and skin was closed using surgical Vetbond (3M). Directly following surgery, 1 mL of PBS was administered subcutaneously to provide post-surgery fluid resuscitation and flunixin meglumine (Phoenix) was administered for post-operative analgesia. Sham mice underwent an identical laparotomy surgical procedure, excluding CLP.

### Bacterial load quantification

Bacterial load was determined by using qPCR targeting the V4 region of the 16s rRNA gene as described elsewhere [Wang et al Cell Gdf15 paper]. Briefly, a correlation curve was built by measuring OD600 of an overnight *E.coli* culture and spread plating colony forming unit (CFU) counts on LB agar plates. Next, this curve was used to build a standard curve with known values for the range of 10e2-10e9 CFU/mL. Genomic DNA was extracted from these colonies and from the organs with DNeasy UltraClearn microbial kit (QIAGEN, 10196-4). After performing qPCR, the values were interpolated according to the standard curve.

### Lipopolysaccharide (LPS)-induced sepsis

Lipopolysaccharide from *E. coli* O55:B5 (Sigma L2880) was freshly resuspended in H2O, prepared in PBS (200 uL), and injected intraperitoneally into mice 2 hour after food removal. The effects of LPS vary significantly from lot to lot. Thus, LD50 was determined for individual lots. LD50 was also determined across the different genotypes, ages, and sexes of mice. In our experiments, the dose varied from 6.5 mg/kg to 15 mg/kg to achieve 50% mortality.

### Sepsis severity evaluation

Clinical signs of sepsis were evaluated and used for scoring disease severity. Clinical scores were assessed by ascending morbidity scale(Berton et al., 2022). Grooming: 0, (Normal); 1, fur that has lost shine/become matte (Dusty); 2, fur becomes erect or bristling (Ruffled). Mobility: 0, mobile without stimulation (Normal); 1, mice are less responsive/mobile to stimuli (Reduced); 2, mice are unresponsive to stimuli (Immobile). Body Position: 0, body is fully extended (Normal); 1, back is arched/curved (Hunched); 2, laying on side at rest (On side). Weight loss: 0, Due to minimal weight loss that occurs in sham control mice, weight loss has been adjusted to allow for surgery-specific weight loss to be mitigated (<10%); 1, moderate weight loss (10-15%); 2, severe weight loss (>15%). After giving one score for each category, the sum of all categories indicates disease score. Importantly, dead mice are given highest score (8) on the day of death, and thereafter removed from scoring. Healthy scores range from 0-2; moderate disease scores range from 3-5; and severe disease scores range from 6-8.

### Cell culture and treatments

#### Primary hepatocyte isolation

Primary hepatocytes were isolated from mice using the collagenase type X perfusion method. Briefly, primary hepatocytes were isolated by collagenase type X (Wako Pure Chemical Industries, ltd, Japan; 039-17864) perfusion method. Cells were washed with hepatocyte wash medium (Thermo Fisher Scientific, Waltham, MA; 17704024), purified by 30% Percoll (GE Healthcare, Chicago, IL; 17089101) density gradient separation, and resuspended in William’s E medium (Thermo Fisher Scientific; 12551032) with 5% fetal bovine serum, 10 nM dexamethasone, and 20 nM insulin. Cells were then seeded on collagen-coated plates at a final density of 3.5 × 10 ^4^ cells/cm. After 4 hours, attached cells were cultured with fresh medium and transduced with the indicated adenoviruses.

For adenovirus transduction, 3 MOI of indicated constructs of adenovirus were added to isolated primary hepatocytes. To measure ER homeostasis, isolated primary hepatocytes were seeded to pre-coated rat tail collagen I (Corning, Cat 370 No. 354236) 24-well plates. After overnight incubation, cells were transfected with the 1.0 µg/well UPRE-luciferase reporter with 0.15 µg/well Renilla luciferase vector (Promega, Cat No. 372 E2261) using polyethylenimine (PEI, Polysciences INC, Cat No. 23966). At 48 hrs post-transfection, activities of Cypridina noctiluca (Clu) luciferase in medium were measured using 1 mM Cypridina cypridina 374 Luciferin (Prolume, Cat No. #305), and the Gaussia luciferase (Gluc) was used as a secretion control measured using 10 mM native coelenterazine (CTZ) (Prolume, Cat No. #303). In the UPRE reporter assay, firefly luciferase and Renilla luciferase were measured using the Dual-Glo Luciferase Assay (Promega, Cat 377 No. E2920). The luminescence was read on a microplate reader (SpectraMax, Molecular Devices, San Jose, 378 CA).

#### HL-1 cell culture and experimental manipulations

HL-1 cell line was purchased from Millipore Sigma (SCC065) and cultured in Claycomb medium (Sigma, 51800C) with other additives as recommended. Cells were cultured on fibronectin (Sigma, F0895)/gelatin (Sigma, G1393). For measuring intracellular calcium, HL-1 cells were seeded in a pre-coated 96-well black plate with optical bottom. Prior to adding appropriate treatments, the cells were extensively washed with Claycomb medium with a multichannel pipette. Rhod-2AM (ThermoFisher Scientific, R1244) was used at 5 µM (30 minutes) with subsequent wash with Claycomb medium. A microplate reader (CLARIOstar PLUS, BMG Labtech, Germany) was used to detect fluorescent signal from the bottom.

### AAV8- and adenovirus-mediated gene delivery *in-vivo*

AAV2/8-TBG-iCre was purchased from Vector Biolabs (titer: 3.0x10^13^ GC/mL). The virus was diluted in saline and 100 µL of 3.0x10^11^ GC/mL was injected into each mouse retro-orbitally. Subsequent experiments were carried out 8 weeks after the injection. Adenovirus delivery was achieved by injecting 1.7x10^10^ ifu in 100 µL of saline retro-orbitally. Subsequent experiments were carried out 2 weeks after the injection.

### Immunohistochemistry

#### Frozen organs

Freshly frozen heart or liver was sectioned into ∼8-10 um slices at -19C, placed on charged glass slides, and kept on dry ice or at -80C until further processing. The sectioned specimen was then fixed in 4% paraformaldehyde (PFA) at room temperature for 15 min, washed with phosphate buffer solution (PBS) 3 times, blocked for 1 hour at room temperature (in PBS with 10% goat serum, 0.1% Triton-X 100, and 1 mM CaCl2) and incubated with primary antibodies (1:300 in PBS with 0.1% goat serum, 0.1% Triton-X 100, and 1 mM CaCl2) overnight at 4C. Next morning, the slides were washed with PBS 3 times, incubated with a secondary antibody conjugated to a fluorophore (1:300 in PBS with 0.1% goat serum, 0.1% Triton-X 100, and 1 mM CaCl2). The slides were then washed with PBS 3 times, mounted with DAPI (Sigma F6057), and imaged using Zeiss 700 (Germany).

### Primary cells

After indicated treatments, cells were washed with PBS 3 times, fixed with 1% PFA for 15 min at room temperature. After this point, the protocol of blocking and staining was similar to that in frozen organs. *Bodipy staining*: freshly frozen heart or liver was sectioned into 8-10 um slices at -19C, placed on charged glass slides, and kept on dry ice or at -80C until further processing. The sectioned specimen was then fixed in 4% paraformaldehyde (PFA) at room temperature for 15 min, washed with PBS 3 times, permeabilized with PBS-T (Triton-X 100 0.1%) for 15 min at room temperature, and incubated with glycine (0.3 M, Sigma W328712) for 15 min at room temperature. Bodipy^TM^ 493/503 (ThermoFisher D3922) was prepared in PBS-T at 2 µM and incubated for 15 min at room temperature. The slides were washed with PBS 3 times, mounted with DAPI, and images were acquired with Zeiss 700 (Germany).

### VLDL isolation

VLDLs were isolated as previously described (Li et al., 2018). Briefly, 5 age-matched male mice were injected with LPS (10 mg/kg) for 2 hours. The blood was then collected from the suprahepatic inferior vena cava and collected into EDTA-treated tubes. The samples were pooled and centrifuged at 2,000*g for 10 min at 4℃, followed by another centrifugation at 15,000*g for 30 min at 4℃. Density was adjusted to 1.063 g/mL by adding appropriate amount of KBr to the plasma on ice. It was then transferred into a propylene centrifuge tube and total volume was adjusted to ∼3.5 mL by adding PBS (density was adjusted to 1.063 g/mL with KBr). The samples were then centrifuged in a fixed-bucket rotor at 100,000*g for 24 hours at 4℃. The top 1/5 of the sample was then transferred into a new tube and stored at 4℃. The VLDL fraction was used for HL-1 experiments on the same day.

### Quantitative RT-PCR (qRT-PCR)

Total RNA was isolated from organs and tissues with Trizol reagent (ThermoFisher 15596026). RNA concentration was determined with a nanodrop and adjusted to 1 ug for RT reaction (ThermoFisher 4368814). 8 ng of cDNA was used for the SYBR green reagent (BioRad 1725270) reaction for qPCR analysis (BioRad). The following are the primers that were used in this study: *16s* FWD: 5’-GTGCCAGCMGCCGCGGTAA-3’, REV:5’-GGACTACHVGGGTWTCTAAT-3’; *Apob* FWD: 5’-ATGGCGTTACGGCAACAAACA-3’, REV:5’CTCCAGATCGTCTTTAGGAAGAGGTTG- 3’*; Ern1* FWD: 5’-GTGACCGAATAGAAAAGGAGGC-3’, REV: 5’-TTCTCCCGCCAGTCCATCTT-3’; *Gapdh* FWD: 5ʹ-TGTGTCCGTCGTGGATCTGA-3ʹ, REV: 5ʹ-CTGCTTCACCACCTTCTTGA T-3ʹ*; Il-1b* FWD: 5’-GCAACTGTTCCTGAACTCAACT-3’, REV: 5’-ATCTTTTGGGGTCCGTC AACT-3’*; Il-6* FWD: 5’-CGCTATGAAGTTCCTCTCTGC-3’, REV: 5’CTAGGTTTGCCGAGTAG ATCTC-3’*; Tnfa* FWD: 5’-CCCTCACACTCAGATCATCTTCT-3’, REV: 5’-GCTACGACGTG GGCTACAG-3’.

### Nuclear fractionation

∼150 mg of freshly harvested liver was homogenized in 1 mL of hypotonic buffer (0.25 M sucrose, 0.02 M HEPES pH 7.5, 0.01 M KCl, 0.0015 M MgCl2, 0.001 M EDTA, 0.001 M EGTA, and protease inhibitors) using mechanical homogenization. The lysate was then filtered through a 100 µm cell strainer and centrifuged at 1,734*g for 20 min at 4℃. The supernatant was further spun at 10,000*g for 30 min at 4℃. The pellet was resuspended in 500 µL of nuclear extraction buffer (0.02 M HEPES pH 7.9, 0.0015 M MgCl2, 0.5 M NaCl, 0.0002 M, 20% glycerol, and protease inhibitors). At this point, the resuspended pellet was passed through a 32G needle until the lysate became clear. Finally, the lysate was spun at 17,968*g for 15 min at 4℃ and briefly sonicated on ice.

### Western blot analysis

Organs and tissues were homogenized in a lysis buffer (50 mM Tris-HCl pH 7.5, 2 mM EDTA, 5 mM EGTA, 30 mM NaF, 40 mM β-glycerophosphate, 10 mM sodium pyrophosphate, 0.3% NP-40) and freshly added protease inhibitor cocktail (Sigma-Aldrich, P8340). Protein concentration was determined by Pierce BCA kit (Thermo Fisher Scientific, 23225) and adjusted samples were subjected to SDS-polyacrylamide gel electrophoresis followed by an overnight transfer onto PVDF membranes (Bio-Rad Laboratories, 1620177). Membranes were immunoblotted with antibodies overnight at 4℃ as follows: IRE1ɑ (Cell Signaling Technology, 14C10), ACTB (Abcam, ab8227), ApoB (Abcam, ab20737), sXBP1 (Biolegend, 647501), Ly-6G (ThermoFisher Scientific, 11-9968-82), cleaved caspase-3 (Cell Signaling Technology, Asp175), AIF1 (ThermoFisher Scientific, BS-1363R). Secondary antibodies were horseradish peroxidase-conjugated goat anti-mouse-IgG (Santa Cruz Biotechnology, sc-2005) and horseradish peroxidase-conjugated anti-rabbit-IgG (CST 7074) at 1:5,000 dilution for 1 hour at RT. The signal was detected using the ChemiDoc Touch Imaging System (Bio-Rad Laboratories), and densitometry analyses of western blot images were carried out with the Image Lab software (Bio-Rad Laboratories).

### High resolution respirometry in cardiac fibers (Oroboros)

High-resolution oxygen consumption was carried using the OROBOROS Oxygraph-2k (O2k, Oroboros instruments, Innsbruck, Austria). Freshly resected left ventricular tissue (∼2.5-4 mg per sample) was pulled into fibers and permeabilized in the presence of saponin (50 µg/mL, Sigma, 47036) 30 min at 4^0^C in BIOPS buffer containing the following: 7.23 mM K2EGTA, 2.77 mM CaK2EGTA, 20 mM imidazole, 0.5 mM DTT, 20 mM Taurine, 5.7 mM ATP, 14.3 mM PCr, 6.56 mM MgCl2-6H2O, 50 mM MES, pH adjusted to 7.1 with 5 N KOH (reagents purchased from Sigma). The digested fibers were then incubated for 10 min at 4^0^C in buffer Z’ containing the following: 105 mM K-MES, 30 mM KCl, 10 mM KH2PO4, 5 mM MgCl2-6H2O, 2.5 mg/mL BSA-FA-free, pH adjusted to 7.4, and 1 mM EGTA was added prior to the experiment (reagents were purchased from Sigma). Measurements were recorded in the presence of 2.5 mM ADP (state 3 respiration, Sigma, A2754) and the absence of ADP (state 2 respiration) while providing palmitoyl-carnitine (0.075 mM, Sigma, 61251) and malate (1 mM, Sigma, 1613881). MCC950 (5 µM) or MitoTEMPO (50 µM, Sigma, SML0737) were added to the fibers during the saponin incubation as well as the subsequent incubation in buffer Z’ (total of 1 hour).

### Blood work

Multiplex cytokine analysis was performed via ThermoFisher Scientific’s ProcartaPlex 7-Plex Panel according to manufacturer’s instructions for plasma cytokine analysis. Samples were analyzed on a Bio-Rad Laboratories Bio-Plex (Luminex 200) analyzer at the University of Iowa Flow Cytometry Core Facility.

Measurement of other parameters, including CTNI (UCP Biosciences, CARD-004A), ALT (Sigma-Aldrich, MAK052), AST (Sigma-Aldrich, MAK055), IL-1b (Biolegend, 432601), TNF-a (Biolegend, 430904), and IL-6 (Biolegend, 431304) according to the manufacturer’s instructions.

### Echocardiography

Transthoracic echocardiograms were performed on conscious mice in the University of Iowa Cardiology Animal Phenotyping Core Laboratory, using a Vevo 2100 Imager (Visual Sonics), as described previously (Hammond et al., 2016; Penny and Hammond, 2017). The echocardiographer was blinded to the treatments and genetic backgrounds of the animals. 2D imaging and M-mode echocardiography were performed in the left ventricle (LV) short- and long-axis planes. Left ventricular end-diastolic volume (LVEDV), end-systolic volume (LVESV), and ejection fractions (LVEF) were calculated with the area-length method.

### LC/MS metabolomics

Data were acquired from an Agilent 6545 UHD QTOF interfaced with an Agilent 1290 UHPLC. Metabolites were separated by using a Millipore Sigma SeQuant ZIC-pHILIC (150 mm x 2.1 mm, 5 μm) column. Solvents were A, 95% water in acetonitrile with 10 mM ammonium acetate and 5 μM phosphate, and B, 100% acetonitrile. A flow rate of 200 μL/min was applied with the following gradient (minutes, %B): 0, 94.7%; 2, 94.7%; 27, 36.8%; 35, 20.0%; 37, 20.0%; 39, 36.8%. For all experiments, 2 μL of metabolic extract was injected. MS parameters were as follows: gas, 200°C 4 L/min; nebulizer, 44 psi; sheath gas, 300°C 12 L/min; capillary, 3kV; fragmentor, 100V; scan rate, one scan/s. MS detection was carried out in both positive and negative modes with a mass range of 65–1,700 Da. Identifications were established by comparing the retention times and fragmentation data of compounds to model standards. All raw data files were converted into mzXML files by using msconvert. Data analysis was performed by using either Agilent’s Profinder or in-house R packages.

### LC/MS Lipidomics

#### Chemicals

LC-MS-grade solvents and mobile phase modifiers were obtained from Honeywell Burdick & Jackson, Morristown, NJ (acetonitrile, isopropanol, formic acid), Fisher Scientific, Waltham, MA (methyl *tert*-butyl ether) and Sigma–Aldrich/Fluka, St. Louis, MO (ammonium formate, ammonium acetate). Lipid standards were obtained from Avanti Polar Lipids, Alabaster, AL (Avanti Mouse SPLASH LIPIDOMIX (330710)) and Cayman Chemical, Ann Arbor, MI, (Palmitic acid-d31 (16497)).

#### Plasma Extraction

Lipid extraction was performed as described previously (Alshehry et al., 2015). In brief, 60 µL of plasma was mixed with 600 µL of butanol:methanol (1:1) which contained a mixture of internal standards (Avanti Mouse Splash, 20 µL per sample; and cholesterol-d7 (100 ug/mL), 10 µL per sample). Samples were vortexed thoroughly and set in a sonicator bath for 1 hour maintained at room temperature. Samples were then centrifuged (14,000xg, 10 min, 20 °C) before transferring the into sample vials for analysis.

#### Liver Extraction

Extraction of lipids was carried out using a biphasic solvent system of cold methanol, methyl tert-butyl ether (MTBE), and water as described by Matyash et al. (J Lipid Res 49(5) (2008) 1137-1146) with some modifications. In a randomized sequence, tissue lipids were extracted in bead-mill tubes (ceramic 1.4 mm, Mo-Bio, Qiagen, Germantown, MD) containing a solution of 225 µL MeOH, 750 µL methyl tert-butyl ether (MTBE), and internal standards. Samples were homogenized in one 30 sec cycle then rested on ice for 1 hr with occasional vortexing. Then 188 µL of PBS was added followed by a brief vortex. Samples were then centrifuged at 14,000 x g for 10 minutes at 4 °C, and the upper phases were collected. Another aliquot of 1 mL MTBE/MeOH/water (10/3/2.5, v/v/v) was added to the bottom aqueous layer followed by a brief vortex. Samples were then centrifuged at 14,000 x g for 10 minutes at 4 °C, the upper phases were combined and evaporated to dryness under speedvac. Lipid extracts were reconstituted in IPA/ACN/water (4:1:1, v/v/v). Concurrently, a process blank sample was prepared and then a pooled quality control (QC) sample was prepared.

#### LC-MS Analysis

Lipid extracts are separated on a Waters Acquity UPLC CSH C18 1.7 µm 2.1 x 100 mm column maintained at 65 °C connected to an Agilent HiP 1290 Sampler, Agilent 1290 Infinity pump, equipped with an Agilent 1290 Flex Cube and Agilent 6545 Accurate Mass Q-TOF dual AJS-ESI mass spectrometer. For positive mode, the source gas temperature is set to 225 °C, with a drying gas flow of 11 L/min, nebulizer pressure of 40 psig, sheath gas temp of 350 °C and sheath gas flow of 11 L/min. VCap voltage is set at 3500 V, nozzle voltage 500V, fragmentor at 110 V, skimmer at 85 V and octopole RF peak at 750 V. For negative mode, the source gas temperature is set to 300 °C, with a drying gas flow of 11 L/min, a nebulizer pressure of 30 psig, sheath gas temp of 350 °C and sheath gas flow 11 L/min. VCap voltage is set at 3500 V, nozzle voltage 75 V, fragmentor at 175 V, skimmer at 75 V and octopole RF peak at 750 V. Samples are analyzed in a randomized order in both positive and negative ionization modes in separate experiments acquiring with the scan range m/z 100 – 1700. Mobile phase A consists of ACN:H2O (60:40 *v/v*) in 10 mM ammonium formate and 0.1% formic acid, and mobile phase B consists of IPA:ACN:H2O (90:9:1 *v/v/v*) in 10 mM ammonium formate and 0.1% formic acid. The chromatography gradient for both positive and negative modes starts at 15% mobile phase B then increases to 30% B over 2.4 min, it then increases to 48% B from 2.4 – 3.0 min, then increases to 82% B from 3 – 13.2 min, then increases to 99% B from 13.2 – 13.8 min where it’s help until 16.7 min and then returned to the initial conditioned and equilibrated for 5 min. Flow is 0.4 mL/min throughout, injection volume is 7 µL for positive and 12 µL negative mode, and tandem mass spectrometry is conducted using the same LC gradient at collision energies of 20 V and 27.5 V, respectively.

### Analysis of Mass Spectrometry Data

QC samples (n=8) and blanks (n=4) are injected throughout the sample queue and ensure the reliability of acquired lipidomics data. Results from LC-MS experiments are collected using Agilent Mass Hunter (MH) Workstation and analyzed using the software packages MH Qual, MH Quant, and Lipid Annotator (Agilent Technologies, Inc.). Results from the positive and negative ionization modes from Lipid Annotator are merged based on the class of lipid identified. For example, we detect PC in both POS and NEG modes, but only use the NEG mode data to tabulate data for those targets. The data table exported from MHQuant is evaluated using Excel where initial lipid targets are parsed based on the following criteria. Only lipids with relative standard deviations (RSD) less than 30% in QC samples and are used for data analysis. Additionally, only lipids with background AUC counts in process blanks that are less than 30% of QC are used for data analysis. The parsed excel data tables are normalized to plasma volume.

### LC/MS lipidomics in liver samples

#### Chemicals

LC-MS-grade solvents and mobile phase modifiers were obtained from Honeywell Burdick & Jackson, Morristown, NJ (acetonitrile, isopropanol, formic acid), Fisher Scientific, Waltham, MA (methyl *tert*-butyl ether) and Sigma–Aldrich/Fluka, St. Louis, MO (ammonium formate, ammonium acetate). Lipid standards were obtained from Avanti Polar Lipids, Alabaster, AL (Avanti Mouse SPLASH LIPIDOMIX (330710)) and Cayman Chemical, Ann Arbor, MI, (Palmitic acid-d31 (16497)).

#### Sample Preparation

Extraction of lipids was carried out using a biphasic solvent system of cold methanol, methyl *tert*-butyl ether (MTBE), and PBS/water (See Ref below) with some modifications. Samples were transferred into ceramic bead mill tubes for homogenization. Samples were each homogenized in 225 µL MeOH with internal standards and 188 µL PBS. Homogenates were collected and added to 750 µL MTBE. Samples were then incubated on ice with occasional vortexing for 1 hr and centrifuged at 15,000 x g for 10 minutes at 4 °C. The organic (upper) layer was collected, and the aqueous (lower) layer was re-extracted with 1 mL of 10:3:2.5 (*v/v/v*) MTBE/MeOH/dd-H2O, briefly vortexed, incubated at RT, and centrifuged at 15,000 x g for 10 minutes at 4 °C. Upper phases were combined and evaporated to dryness under speedvac. Lipid extracts were reconstituted in 750 µL of IPA/ACN/water (4:1:1) and transferred to an LC-MS vial for analysis. Concurrently, a process blank sample was prepared and a pooled quality control (QC) sample was prepared by taking equal volumes from each sample after final resuspension. Extraction protocol based on (Matyash et al., 2008).

#### LC-MS

Lipid extracts were separated on an Acquity UPLC CSH C18 column (2.1 x 100 mm; 1.7 µm) coupled to an Acquity UPLC CSH C18 VanGuard precolumn (5 × 2.1 mm; 1.7 µm) (Waters, Milford, MA) maintained at 65 °C connected to an Agilent HiP 1290 Sampler, Agilent 1290 Infinity pump, and Agilent 6545 Accurate Mass Q-TOF dual AJS-ESI mass spectrometer (Agilent Technologies, Santa Clara, CA). Samples were analyzed in a randomized order in both positive and negative ionization modes in separate experiments acquiring with the scan range m/z 100 – 1700. For positive mode, the source gas temperature was set to 225 °C, with a drying gas flow of 11 L/min, nebulizer pressure of 40 psig, sheath gas temp of 350 °C and sheath gas flow of 11 L/min. VCap voltage is set at 3500 V, nozzle voltage 500V, fragmentor at 110 V, skimmer at 85 V and octopole RF peak at 750 V. For negative mode, the source gas temperature was set to 300 °C, with a drying gas flow of 11 L/min, a nebulizer pressure of 30 psig, sheath gas temp of 350 °C and sheath gas flow 11 L/min. VCap voltage was set at 3500 V, nozzle voltage 75 V, fragmentor at 175 V, skimmer at 75 V and octopole RF peak at 750 V. Mobile phase A consisted of ACN:H2O (60:40, *v/v*) in 10 mM ammonium formate and 0.1% formic acid, and mobile phase B consisted of IPA:ACN:H2O (90:9:1, *v/v/v*) in 10 mM ammonium formate and 0.1% formic acid. For negative mode analysis the modifiers were changed to 10 mM ammonium acetate. The chromatography gradient for both positive and negative modes started at 15% mobile phase B then increased to 30% B over 2.4 min, it then increased to 48% B from 2.4 – 3.0 min, then increased to 82% B from 3 – 13.2 min, then increased to 99% B from 13.2 – 13.8 min where it’s held until 16.7 min and then returned to the initial conditions and equilibriated for 5 min. Flow was 0.4 mL/min throughout, with injection volumes of 1 µL for positive and 10 µL negative mode. Tandem mass spectrometry was conducted using iterative exclusion, the same LC gradient at collision energies of 20 V and 27.5 V in positive and negative modes, respectively.

#### Analysis of Mass Spectrometry Data

For data processing, Agilent MassHunter (MH) Workstation and software packages MH Qualitiative and MH Quantitative were used. The pooled QC (n=8) and process blank (n=4) were injected throughout the sample queue to ensure the reliability of acquired lipidomics data. For lipid annotation, accurate mass and MS/MS matching was used with the Agilent Lipid Annotator library and LipidMatch [236]. Results from the positive and negative ionization modes from Lipid Annotator were merged based on the class of lipid identified. Data exported from MH Quantitative was evaluated using Excel where initial lipid targets are parsed based on the following criteria. Only lipids with relative standard deviations (RSD) less than 30% in QC samples are used for data analysis. Additionally, only lipids with background AUC counts in process blanks that are less than 30% of QC are used for data analysis. The parsed excel data tables are normalized based on the ratio to class-specific internal standards, then to tissue mass and sum prior to statistical analysis.

### Statistics

Data were analyzed in GraphPad Prism version 9.3.1. Data are presented as mean ± standard error of the mean (SEM). The sample size was chosen based on the lab’s previous experience and other reports in the field. To ensure normality, the Shapiro-Wilk test was used in RStudio for each dataset, wherein the p-value > 0.05 dictated normally distributed datasets. Following the Shapiro-Wilk test, a parametric test was carried out. Multiple group comparisons were calculated by 2-way ANOVA with Tukey post-hoc test. Pairwise testing was carried out by the two-sided unpaired Student’s t-test.

---

**Supplemental Figure 1.**
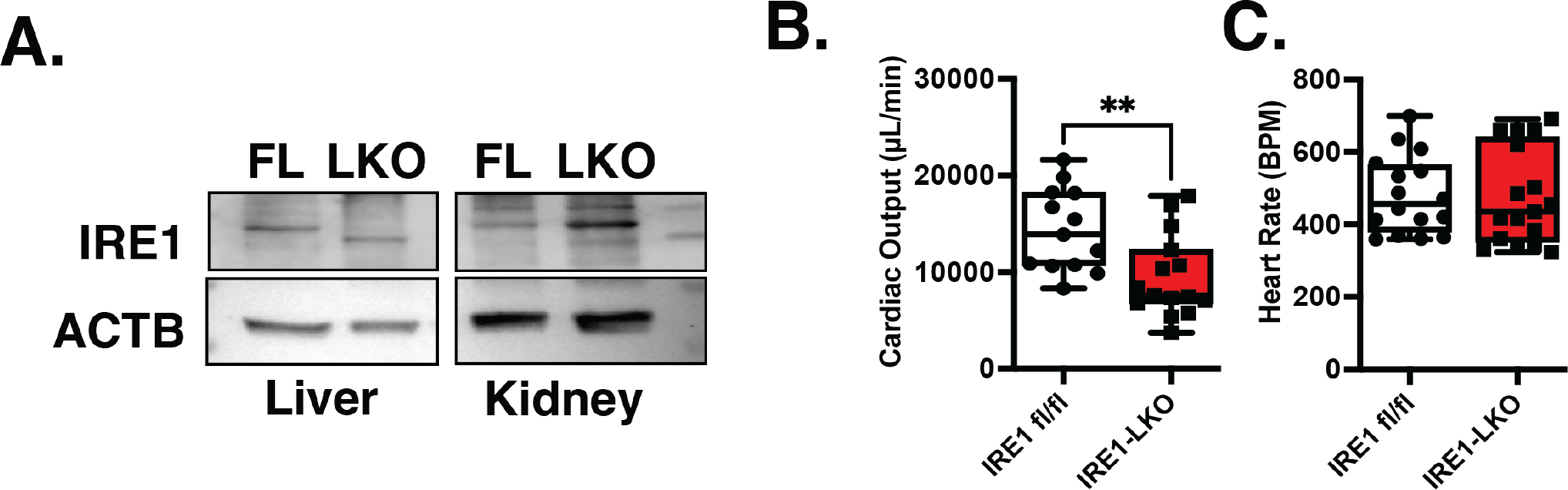
Genetic deletion of IRE1 in the liver and echocardiography parameters. **A.** Western blot analysis of IRE1 and ACTB in the liver and kidney from IRE1^FL/FL^ and IRE1^TBG-CRE^ mice under basal conditions. **B&C.** Cardiac output and heart rate in IRE1^FL/FL^ and IRE1^TBG-CRE^ challenged with 16 hours of LD50 LPS. Data are shown as means ± SEM. *Indicates statistically significant genetic effects. Student’s t-test was used to compare means between groups in (**B&C**).

**Supplemental Figure 2.**
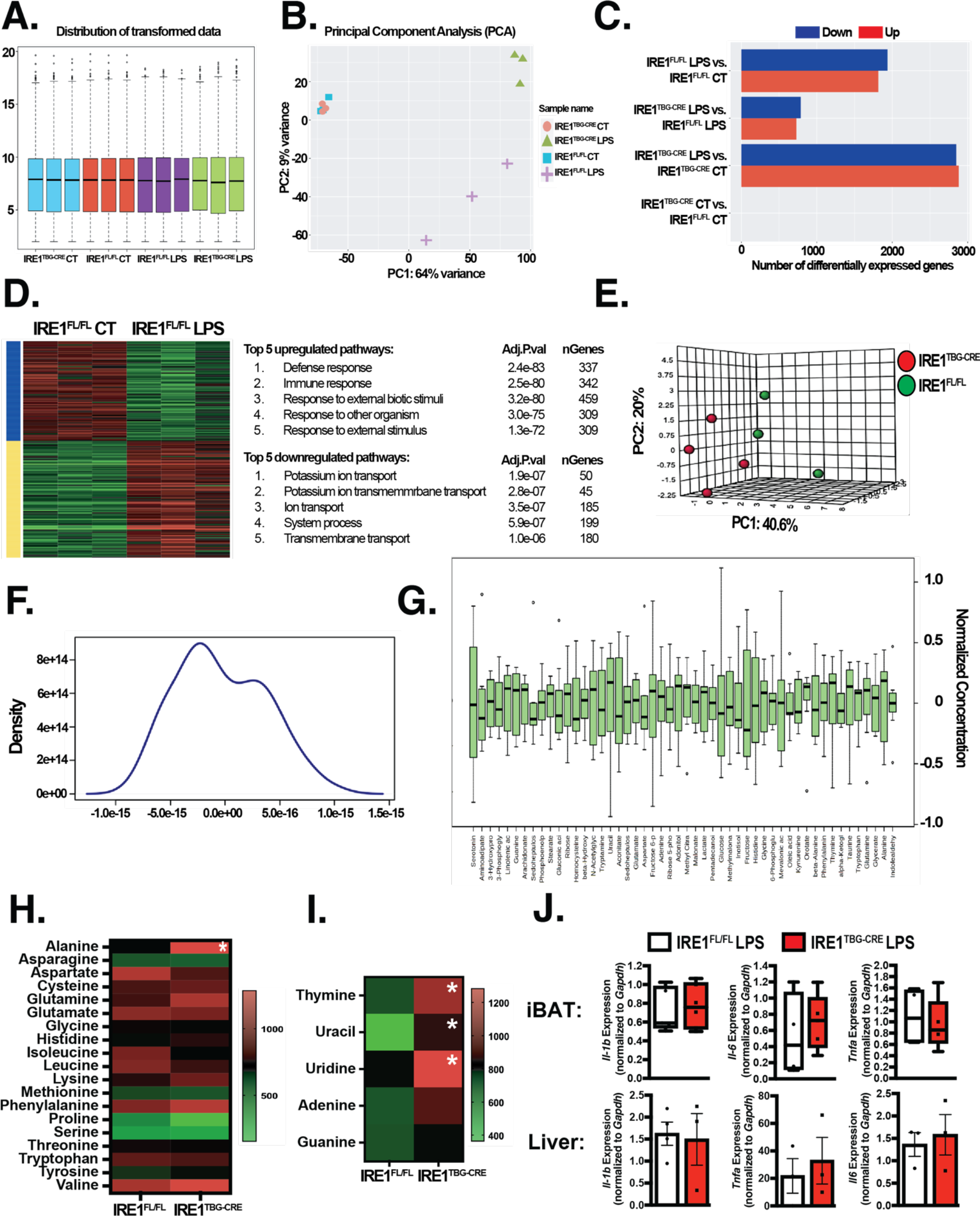
RNA-Seq and GC-MS analyses in LPS-challenged IRE1^FL/FL^ and IRE1^TBG-CRE^ mice. **A-D.** RNA-Seq analysis in hearts from IRE1^FL/FL^ and IRE1^TBG-CRE^ mice after 16 hours of PBS or LPS LD50 challenge; distribution of transformed data in (**A**), principal component analysis in (**B**), up- and down- regulated transcripts in indicated groups in (**C**), and top 5 up- and down-regulated pathways in IRE1^FL/FL^ LPS vs. IRE1^FL/FL^ CT in (**D**); n = 3 age-matched male mice per group; iDEP96 online suite was used for data analysis. **E-I**. GC/MS steady-state metabolomic analysis in hearts from IRE1^FL/FL^ and IRE1^TBG-CRE^ mice after 16 hours of LPS LD50 challenge; principal component analysis in (**E**), density plot of detected metabolites in (**F**), normalized concentration of transformed metabolites in (**G**), heatmap of relative levels of detected amino acids in (**H**), heatmap of relative levels of nucleotides in (**I**); n = 3-4 age-matched male mice. **J**. Top: inflammatory cytokines in iBAT from IRE1^FL/FL^ and IRE1^TBG-CRE^ mice after 16 hours of LPS LD50 challenge measured by qRT-PCR and normalized to *Gapdh;* Bottom: inflammatory cytokines in livers from IRE1^FL/FL^and IRE1^TBG-CRE^ mice after 16 hours of LPS LD50 challenge measured by qRT-PCR and normalized to *Gapdh;* n = 3-4 age-matched male mice. Data are shown as means ± SEM. *Indicates statistically significant genetic effects.

**Supplemental Figure 3.**
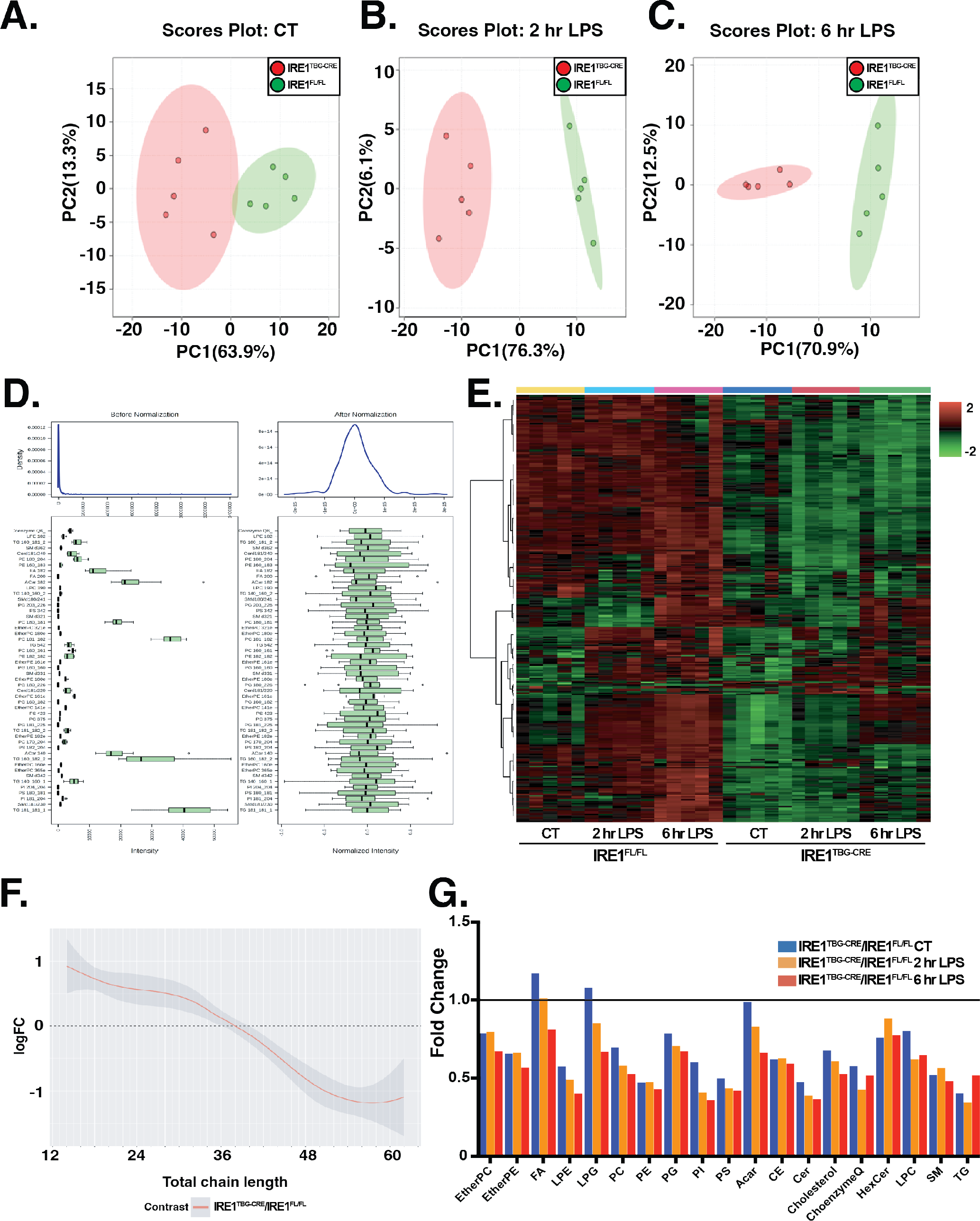
LC/MS lipidomic analysis in IRE1^FL/FL^ and IRE1^TBG-CRE^ mice challenged with LPS. **A-G**. LC/MS lipidomic analysis of plasma isolated from suprahepatic inferior vena cava from IRE1^FL/FL^ and IRE1^TBG-CRE^ mice challenged with LPS at indicated time points; principal component analysis in (**A-C**); normalized intensity of detected lipid species before and after transformation in (**D**), heatmap of detected individual lipid species compared between IRE1^FL/FL^ and IRE1^TBG-CRE^ mice challenged with LPS at indicated time points in (**E**), representation of fold change of total carbon chains in detected lipids between IRE1^FL/FL^and IRE1^TBG-CRE^ mice under basal conditions in (**F**), fold change of lipid groups between IRE1^FL/FL^ and IRE1^TBG-CRE^ mice challenged with LPS at indicated time points in (**G**); n = 5 age-matched male mice.

**Supplemental Figure 4.**
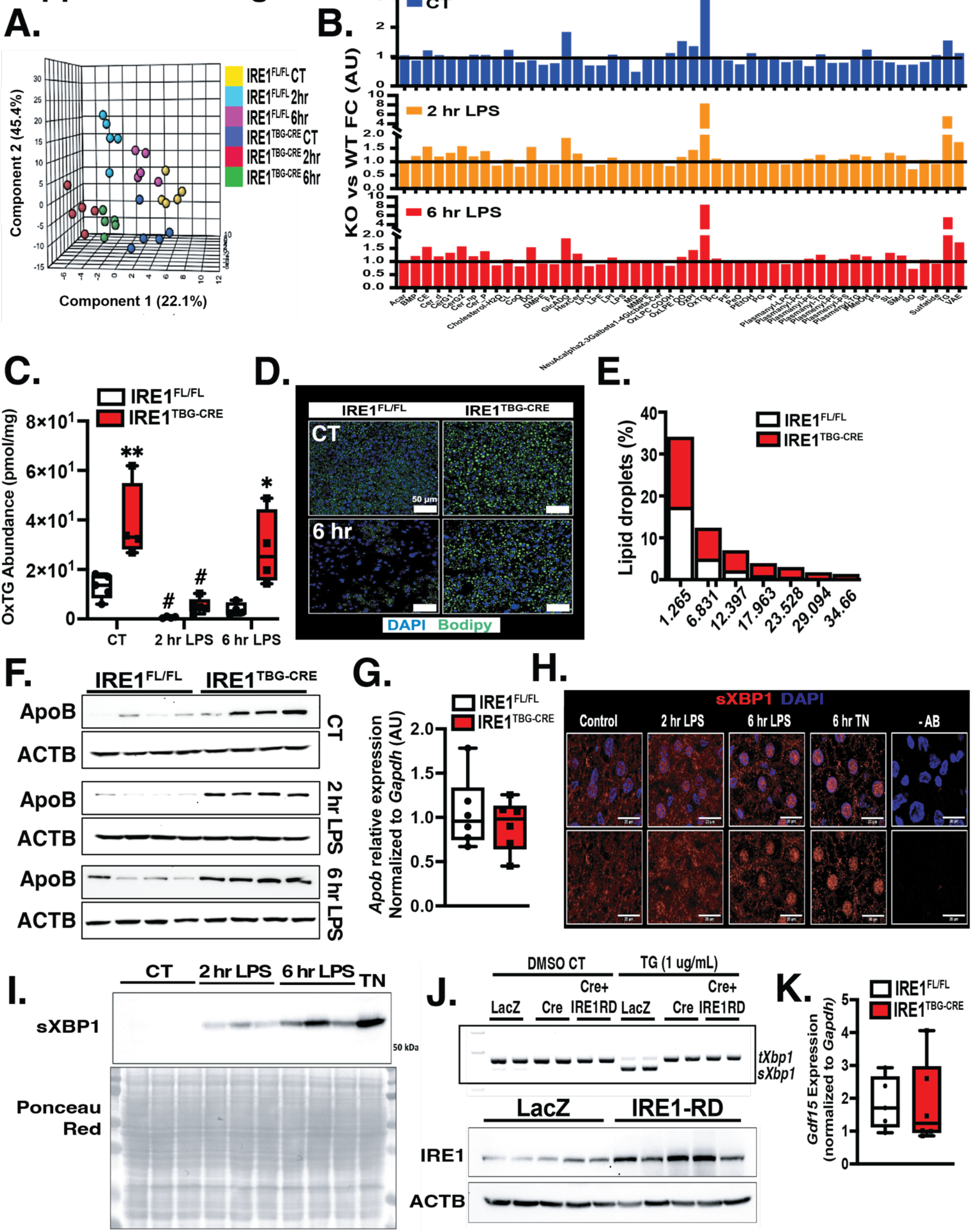
Intrahepatic characterization of lipid landscape in IRE1^FL/FL^ and IRE1^TBG-CRE^ mice challenged with LPS. **A-C**. LC/MS lipidomic analysis in livers from IRE1^FL/FL^ and IRE1^TBG-CRE^ mice challenged with LPS at indicated time points; normalized intensity before and after transformation of data in (**A**). Principal component analysis. (**B**). Fold change of detected lipid groups in IRE1^TBG-CRE^ vs. IRE1^FL/FL^ livers; (**C**). Oxidized TG (OxTG) abundance in the liver; n = 5 age-matched male mice. **D**. Representative confocal images of lipid droplets detected with Bodipy dye in livers from IRE1^FL/FL^ and IRE1^TBG-CRE^ mice under basal conditions and 6 hours after LPS LD50 challenge. **E**. Distribution of lipid droplet sizes in livers from IRE1^FL/FL^ and IRE1^TBG-CRE^ mice under basal conditions; n = 5 age-matched male mice. **F**. Western blot analysis of intrahepatic ApoB from IRE1^FL/FL^ and IRE1^TBG-CRE^ mice challenged with LPS at indicated time points; n = 4 age-matched male mice. **G**. Transcript level of *Apob* from IRE1^FL/FL^ and IRE1^TBG-CRE^ mice under basal conditions measured by qRT-PCR and normalized to *Gapdh;* n = 6 age-matched male mice. **H**. Representative confocal imaging of sXBP1 in livers from WT animals challenged with LPS LD50 or tunicamycin (TN) at indicated time points. **I.** Western blot analysis of sXBP1 and ponceau red in nuclear fractions isolated from livers of IRE1^FL/FL^ and IRE1^TBG-CRE^ mice after LPS or TN challenge for indicated time points; n = 3 age-matched male mice. **J**. Top: Spliced *Xbp1* measured by RT-PCR in primary hepatocytes from IRE1^FL/FL^ mice transduced with Ad-LacZ, Ad-Cre, or Ad-Cre+Ad-IRE1-RD challenged with TN for 2 hours; Bottom: western blot analysis of intrahepatic levels of IRE1 and ACTB in WT mice transduced with adenovirus-LacZ or adenovirus-IRE1-RD for 2 weeks; n = 5 age-matched male mice. K. *Gdf15* transcript expression level in livers from IRE1^FL/FL^ and IRE1^TBG-CRE^ mice under basal conditions measured by qRT-PCR and normalized to *Gapdh*; n = 5-6 age-matched male mice. Data are shown as means ± SEM. *Indicates statistically significant genetic effects and # indicates effects of treatments. Two-way ANOVA followed by Tukey’s multiple comparisons test was used in (**C**). Student’s t-test was used to determine statistical significance in (**G**)&(**K**).

## Notes

### Competing Interest Statement

The authors have declared no competing interest.

