## Supplementary material for "Hepatic IRE1 Protects Against Septic Cardiac Failure": Manuscript

### Supplemental Figure 1

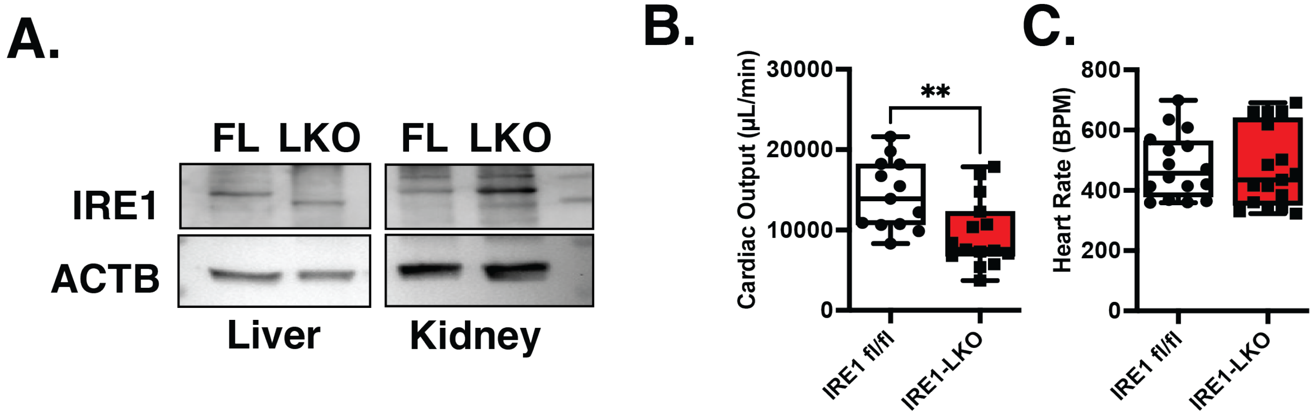


**Supplemental Figure 1. Genetic deletion of IRE1 in the liver and echocardiography parameters.**

**A.** Western blot analysis of IRE1 and ACTB in the liver and kidney from $IRE1^{FL/FL}$ and $IRE1^{TBG\text{-}CRE}$ mice under basal conditions. **B&C.** Cardiac output and heart rate in $IRE1^{FL/FL}$ and $IRE1^{TBG\text{-}CRE}$ challenged with 16 hours of LD50 LPS. Data are shown as means ± SEM. *Indicates statistically significant genetic effects. Student's t-test was used to compare means between groups in (**B&C**).

### Supplemental Figure 2

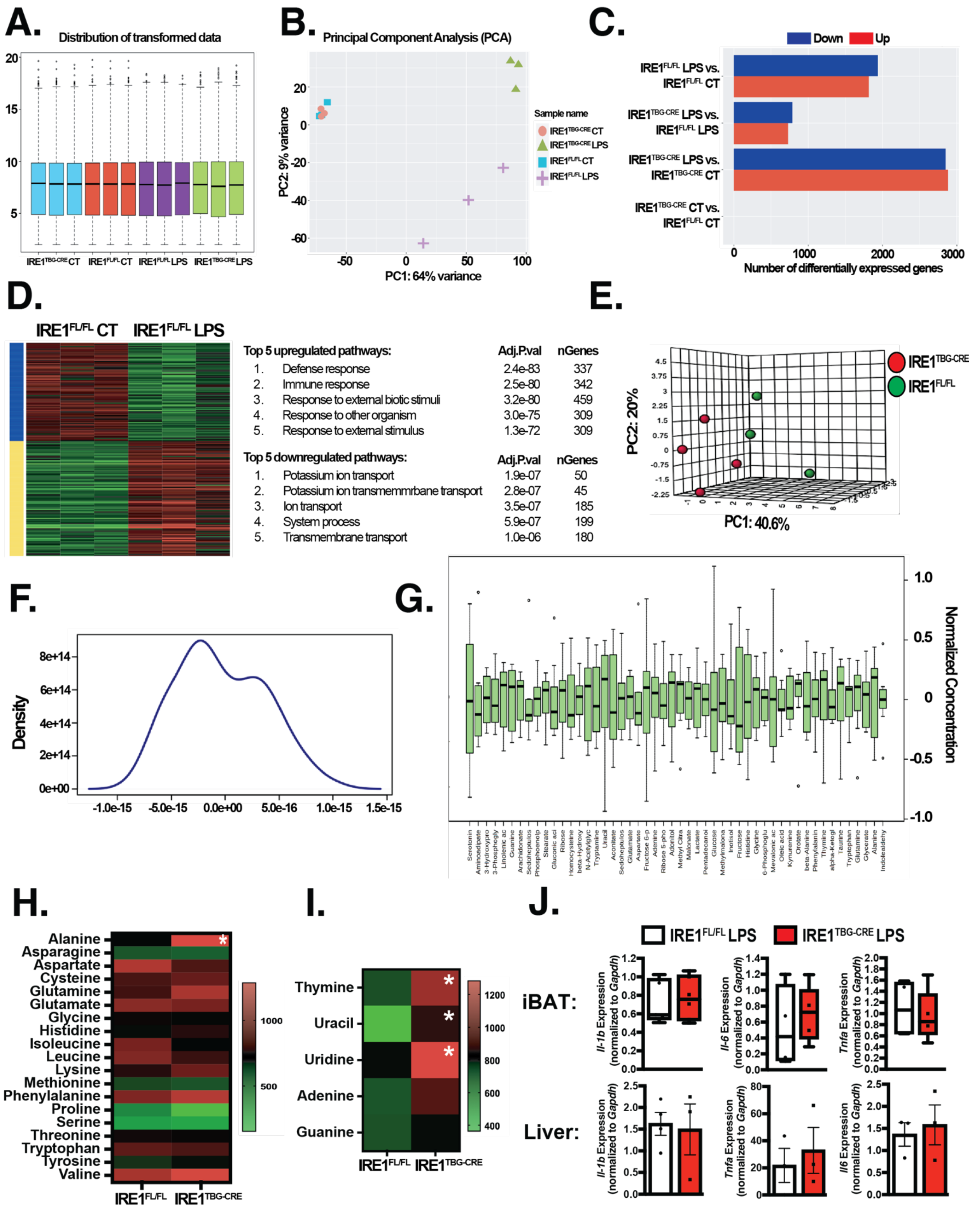

**Supplemental Figure 2. RNA-Seq and GC-MS analyses in LPS-challenged IRE1$^{FL/FL}$ and IRE1$^{TBG-CRE}$ mice.**

**A-D.** RNA-Seq analysis in hearts from IRE1$^{FL/FL}$ and IRE1$^{TBG-CRE}$ mice after 16 hours of PBS or LPS LD50 challenge; distribution of transformed data in (**A**), principal component analysis in (**B**), up- and down-regulated transcripts in indicated groups in (**C**), and top 5 up- and down-regulated pathways in IRE1$^{FL/FL}$ LPS vs. IRE1$^{FL/FL}$ CT in (**D**); n = 3 age-matched male mice per group; iDEP96 online suite was used for data analysis. **E-I**. GC/MS steady-state metabolomic analysis in hearts from IRE1$^{FL/FL}$ and IRE1$^{TBG-CRE}$ mice after 16 hours of LPS LD50 challenge; principal component analysis in (**E**), density plot of detected metabolites in (**F**), normalized concentration of transformed metabolites in (**G**), heatmap of relative levels of detected amino acids in (**H**), heatmap of relative levels of nucleotides in (**I**); n = 3-4 age-matched male mice. **J**. Top: inflammatory cytokines in iBAT from IRE1$^{FL/FL}$ and IRE1$^{TBG-CRE}$ mice after 16 hours of LPS LD50 challenge measured by qRT-PCR and normalized to *Gapdh;* Bottom: inflammatory cytokines in livers from IRE1$^{FL/FL}$ and IRE1$^{TBG-CRE}$ mice after 16 hours of LPS LD50 challenge measured by qRT-PCR and normalized to *Gapdh;* n = 3-4 age-matched male mice. Data are shown as means ± SEM. *Indicates statistically significant genetic effects.

### Supplemental Figure 3

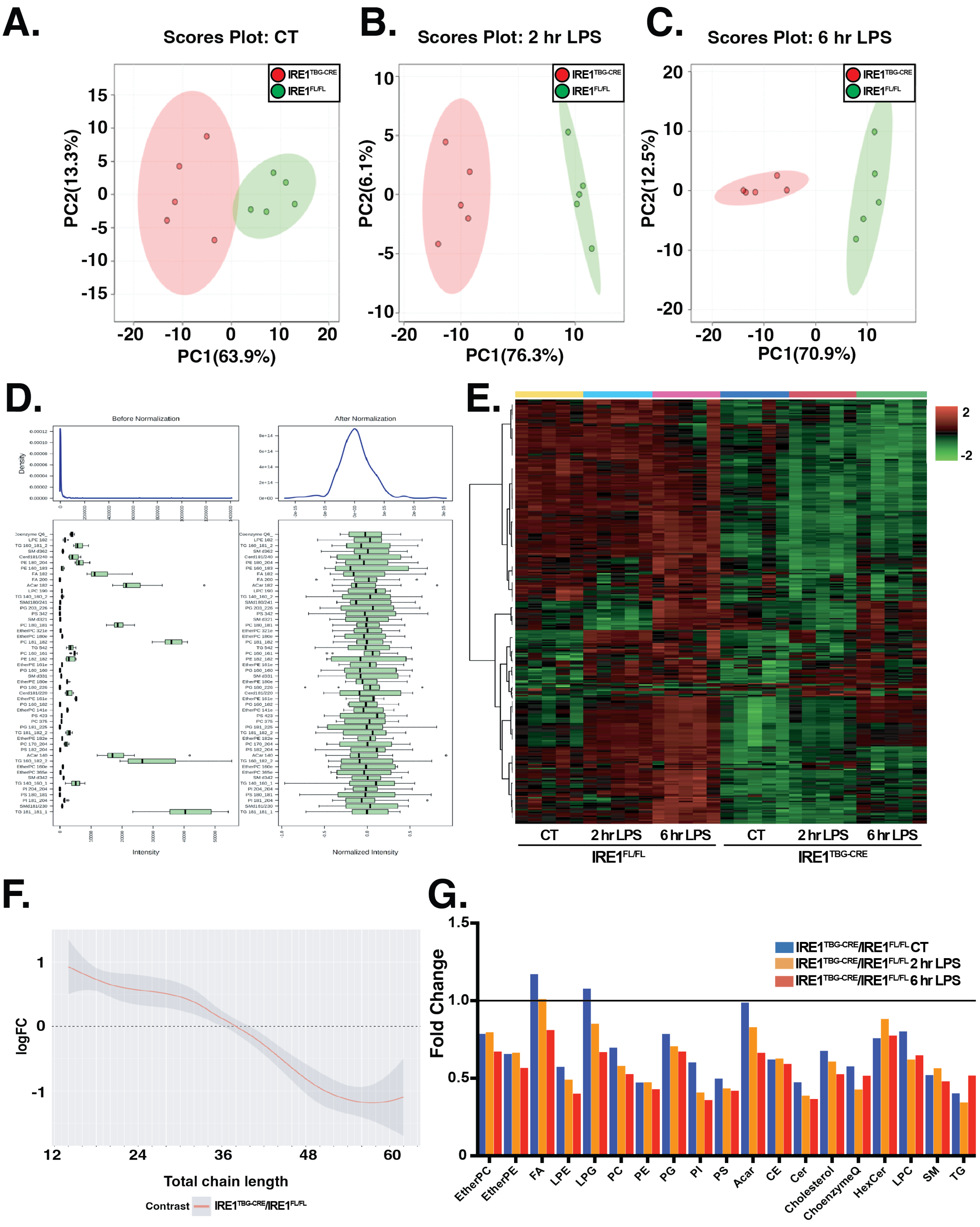

**Supplemental Figure 3. LC/MS lipidomic analysis in IRE1$^{FL/FL}$ and IRE1$^{TBG\text{-}CRE}$ mice challenged with LPS.**

**A-G**. LC/MS lipidomic analysis of plasma isolated from suprahepatic inferior vena cava from IRE1$^{FL/FL}$ and IRE1$^{TBG\text{-}CRE}$ mice challenged with LPS at indicated time points; principal component analysis in (**A-C**); normalized intensity of detected lipid species before and after transformation in (**D**), heatmap of detected individual lipid species compared between IRE1$^{FL/FL}$ and IRE1$^{TBG\text{-}CRE}$ mice challenged with LPS at indicated time points in (**E**), representation of fold change of total carbon chains in detected lipids between IRE1$^{FL/FL}$ and IRE1$^{TBG\text{-}CRE}$ mice under basal conditions in (**F**), fold change of lipid groups between IRE1$^{FL/FL}$ and IRE1$^{TBG\text{-}CRE}$ mice challenged with LPS at indicated time points in (**G**); n = 5 age-matched male mice.

### Supplemental Figure 4

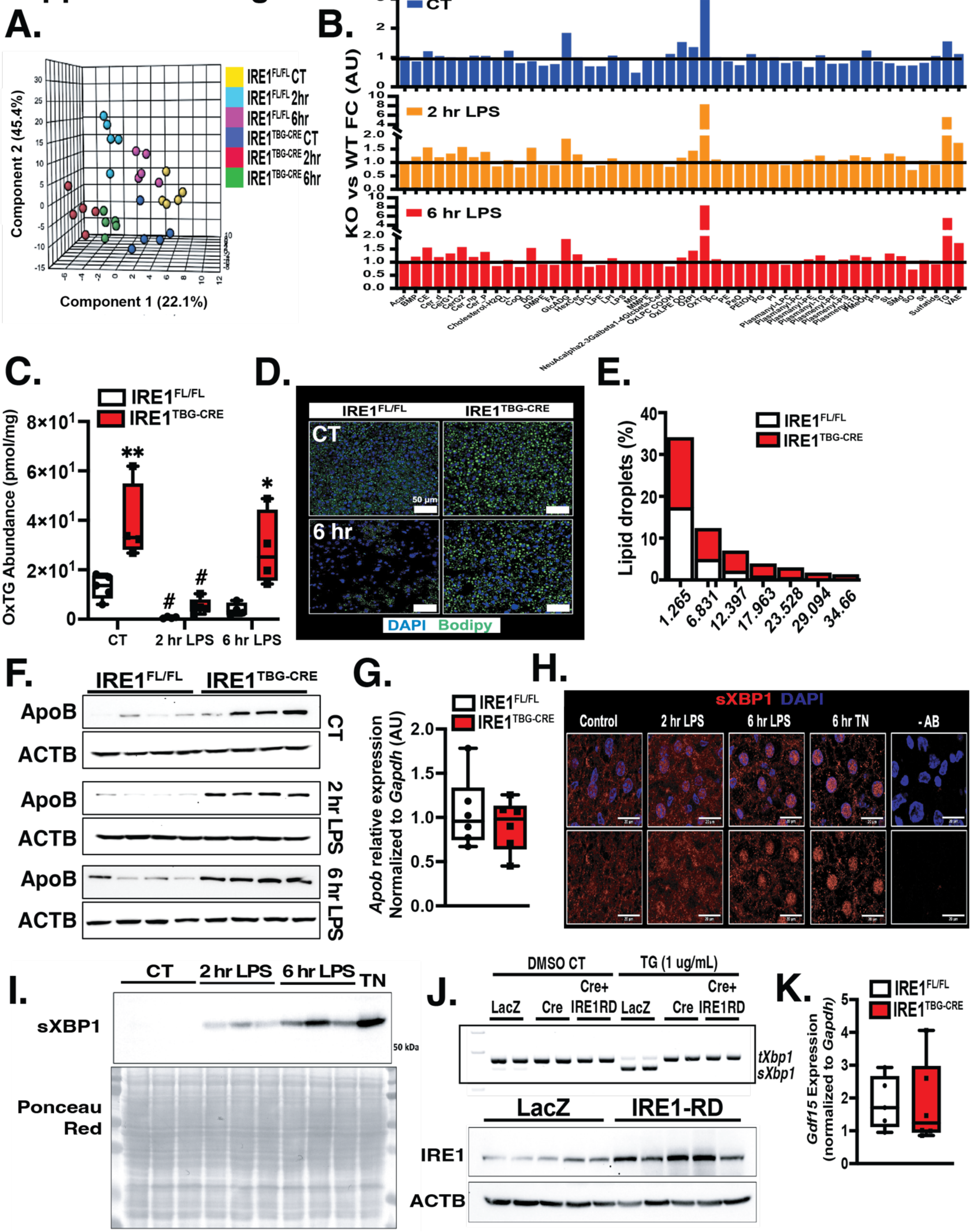

**Supplemental Figure 4. Intrahepatic characterization of lipid landscape in $IRE1^{FL/FL}$ and $IRE1^{TBG-CRE}$ mice challenged with LPS.**

**A-C**. LC/MS lipidomic analysis in livers from $IRE1^{FL/FL}$ and $IRE1^{TBG-CRE}$ mice challenged with LPS at indicated time points; normalized intensity before and after transformation of data in (**A**). Principal component analysis. (**B**). Fold change of detected lipid groups in $IRE1^{TBG-CRE}$ vs. $IRE1^{FL/FL}$ livers; (**C**). Oxidized TG (OxTG) abundance in the liver; n = 5 age-matched male mice. **D**. Representative confocal images of lipid droplets detected with Bodipy dye in livers from $IRE1^{FL/FL}$ and $IRE1^{TBG-CRE}$ mice under basal conditions and 6 hours after LPS LD50 challenge. **E**. Distribution of lipid droplet sizes in livers from $IRE1^{FL/FL}$ and $IRE1^{TBG-CRE}$ mice under basal conditions; n = 5 age-matched male mice. **F**. Western blot analysis of intrahepatic ApoB from $IRE1^{FL/FL}$ and $IRE1^{TBG-CRE}$ mice challenged with LPS at indicated time points; n = 4 age-matched male mice. **G**. Transcript level of *Apob* from $IRE1^{FL/FL}$ and $IRE1^{TBG-CRE}$ mice under basal conditions measured by qRT-PCR and normalized to *Gapdh;* n = 6 age-matched male mice. **H**. Representative confocal imaging of sXBP1 in livers from WT animals challenged with LPS LD50 or tunicamycin (TN) at indicated time points. **I.** Western blot analysis of sXBP1 and ponceau red in nuclear fractions isolated from livers of $IRE1^{FL/FL}$ and $IRE1^{TBG-CRE}$ mice after LPS or TN challenge for indicated time points; n = 3 age-matched male mice. **J**. Top: Spliced *Xbp1* measured by RT-PCR in primary hepatocytes from $IRE1^{FL/FL}$ mice transduced with Ad-LacZ, Ad-Cre, or Ad-Cre+Ad-IRE1-RD challenged with TN for 2 hours; Bottom: western blot analysis of intrahepatic levels of IRE1 and ACTB in WT mice transduced with adenovirus-LacZ or adenovirus-IRE1-RD for 2 weeks; n = 5 age-matched male mice. K. *Gdf15* transcript expression level in livers from $IRE1^{FL/FL}$ and $IRE1^{TBG-CRE}$ mice under basal conditions measured by qRT-PCR and normalized to *Gapdh*; n = 5-6 age-matched male mice. Data are shown as means ± SEM. *Indicates statistically significant genetic effects and # indicates effects of treatments. Two-way ANOVA followed by Tukey's multiple comparisons test was used in (**C**). Student's t-test was used to determine statistical significance in (**G**)&(**K**).
